# An evolutionary paradigm favoring crosstalk between bacterial two-component signaling systems

**DOI:** 10.1101/2022.05.18.492451

**Authors:** Bharadwaj Vemparala, Arjun Valiya Parambathu, Deepak Kumar Saini, Narendra M Dixit

## Abstract

The prevalent paradigm governing bacterial two-component signaling systems (TCSs) is specificity, wherein the histidine kinase (HK) of a TCS exclusively activates its cognate response regulator (RR). Crosstalk, where HKs activate noncognate RRs, is considered evolutionarily disadvantageous because it can compromise adaptive responses by leaking signals. Yet, crosstalk is observed in several bacteria. Here, to resolve this paradox, we propose an alternative paradigm where crosstalk can be advantageous. We envisioned ‘programmed’ environments, wherein signals appear in predefined sequences. In such environments, crosstalk that primes bacteria to upcoming signals may improve adaptive responses and confer evolutionary benefits. To test this hypothesis, we employed mathematical modeling of TCS signaling networks and stochastic evolutionary dynamics simulations. We considered the comprehensive set of bacterial phenotypes, comprising thousands of distinct crosstalk patterns, competing in varied signaling environments. Our simulations predicted that in programmed environments phenotypes with crosstalk facilitating priming would outcompete phenotypes without crosstalk. In environments where signals appear randomly, bacteria without crosstalk would dominate, explaining the specificity widely seen. Additionally, a testable prediction was that the phenotypes selected in programmed environments would display ‘one-way’ crosstalk, ensuring priming to ‘future’ signals. Interestingly, the crosstalk networks we deduced from available data on TCSs of *Mycobacterium tuberculosis* all displayed one-way crosstalk, offering strong support to our predictions. Our study thus identifies potential evolutionary underpinnings of crosstalk in bacterial TCSs, suggests a reconciliation of specificity and crosstalk, makes testable predictions of the nature of crosstalk patterns selected, and has implications for understanding bacterial adaptation and the response to interventions.

**IMPORTANCE:** Bacteria use two-component signaling systems (TCSs) to sense and respond to environmental changes. The prevalent paradigm governing TCSs is specificity, where signal flow through TCSs is insulated; leakage to other TCSs is considered evolutionarily disadvantageous. Yet, crosstalk between TCSs is observed in many bacteria. Here, we present a potential resolution of this paradox. We envision programmed environments, wherein stimuli appear in predefined sequences. Crosstalk that primes bacteria to upcoming stimuli could then confer evolutionary benefits. We demonstrate this benefit using mathematical modeling and evolutionary simulations. Interestingly, we found signatures of predicted crosstalk patterns in *Mycobacterium tuberculosis*. Furthermore, specificity was selected in environments where stimuli occurred randomly, thus reconciling specificity and crosstalk. Implications follow for understanding bacterial evolution and for interventions.

## INTRODUCTION

Bacteria sense and respond to environmental cues predominantly via two-component signaling systems (TCSs) (1). The first component of a TCS is the transmembrane histidine kinase (HK). The HK detects the stimulus, which typically is a chemical ligand, and gets autophosphorylated. The phosphorylated HK (HK-P) binds to and transfers its phosphoryl group to the response regulator (RR), the second component of the TCS. Phosphorylated RR (RR-P) typically dimerizes and triggers changes in downstream gene expression, mounting a response to the stimulus (1, 2). Cognate HK-RR pairs, which belong to a TCS, are generally co-expressed under a single promoter in an operon (3), and are often upregulated as part of the response to the stimulus (1, 2).

Bacteria can have many tens of distinct TCSs, each performing a different function (1). Evolutionary pressure is thought to have rendered TCSs specific: the HK of a TCS rarely phosphorylates the RR of another TCS (4). Crosstalk between TCSs, defined as phosphotransfer from the HK of one TCS to the RR of another TCS, is considered disadvantageous because it dissipates the signal, decreasing the concentration of the cognate RR-P, and thereby weakening the response (4). Moreover, unwanted responses due to gene expression downstream of noncognate RR-Ps might get triggered. Bacteria typically acquire novel TCSs through gene duplication (5), which would naturally allow crosstalk before diversification of the TCSs into distinct pathways (6, 7). Several experimental and modeling studies have argued that despite the extensive homology between TCS proteins, there is strong evolutionary pressure for these paralogs to be specific (5, 8-13). For instance, crosstalk between TCSs can be abrogated by as few as two mutations, indicative of the evolutionary pressure favoring specificity (8). Further, during the evolution of new TCSs post gene duplication, bacteria have been predicted to eliminate crosstalk before new TCS functionalities can arise (9). The sequence space occupied by the paralogs is thought to be sparse, allowing easy establishment of such specificity (12).

Yet, crosstalk between bacterial TCSs continues to be observed, and, in some bacteria, in significant measure. Approximately 3% of the 850 interactions between TCS proteins in *E. coli*, for instance, were between noncognate HK-RR pairs (14). A substantially larger fraction, ∼50% of the 23 interactions, were between noncognate pairs in *M. tuberculosis* (15). Given the evolutionary advantages of specificity together with the relative ease of establishing it, the observed crosstalk is puzzling. Indeed, in some organisms, such as *C. crescentus* (16) and *M. xanthus* (17), no crosstalk has been observed among hundreds of interactions. The observed crosstalk may thus not be attributable to chance and may instead have evolutionary underpinnings. Unraveling potential evolutionary advantages of crosstalk is expected to have important implications for our understanding of bacterial adaptation, survival, and response to interventions (1, 18, 19).

Here, we conceived of an evolutionary paradigm in which crosstalk could be beneficial. We hypothesized that in programmed environments, where signals consistently appear in a predefined sequence, crosstalk between TCSs that would prime the bacterium to upcoming signals might confer an evolutionary advantage. To test this hypothesis, we constructed a mechanistic mathematical model of generalized multi-TCS signaling networks and performed comprehensive evolutionary dynamics simulations. We challenged model predictions with available experimental observations and found evidence in support of our hypothesis. Additionally, we arrived at a plausible synthesis of the seemingly conflicting observations of specificity and crosstalk in bacterial TCS systems.

## RESULTS

### Crosstalk can confer a fitness advantage in programmed environments

We first considered a hypothetical environment involving N=2 signals, denoted I_1_ and I_2_, recognized by two TCSs of a bacterium, TCS_1_ and TCS_2_, made up of the proteins HK_1_ and RR_1_ and HK_2_ and RR_2_, respectively. Depending on the nature of interactions between the TCSs, four phenotypes could exist (Fig. 1a): 1) with no crosstalk (phenotype 1); 2) with crosstalk between HK_1_ and RR_2_ (phenotype 2); 3) with crosstalk between HK_2_ and RR_1_ (phenotype 3); and 4) with bidirectional crosstalk (phenotype 4). We developed a detailed model of signal transduction in a TCS network, allowing for all possible crosstalk patterns between the TCSs (Methods). The model builds on previous models of TCS signaling (9, 15, 20, 21), generalizing them to multi-TCS networks with crosstalk. The novelty of our approach lies in recognizing and incorporating the role of the environment. We applied our model to each of the four phenotypes. We first considered the scenario representing a programmed environment. Specifically, we let the signal I_1_ be followed by I_2_. For simplicity, we let the signals be identical except for the time of their onset (Fig. 1b). We also assumed the signals to be square pulses arriving in quick succession, mimicking the typical way environments impose stresses (22); we considered alternative signal types below. Using the model, we predicted the concentrations of RR_1_-P and RR_2_-P over time (Fig. 1b top panel) as a proxy for the responses of the bacteria to the two stimuli. Further, we estimated the fitness, *ϕ*_1_ and *ϕ*_2_, of the bacteria associated with the responses of the two TCSs, and the overall fitness, ⟨*ϕ*⟩, combining the two (Fig. 1b bottom panel). The fitness was determined by the strength of the cognate responses to the individual stimuli (Methods).

**Fig. 1.**
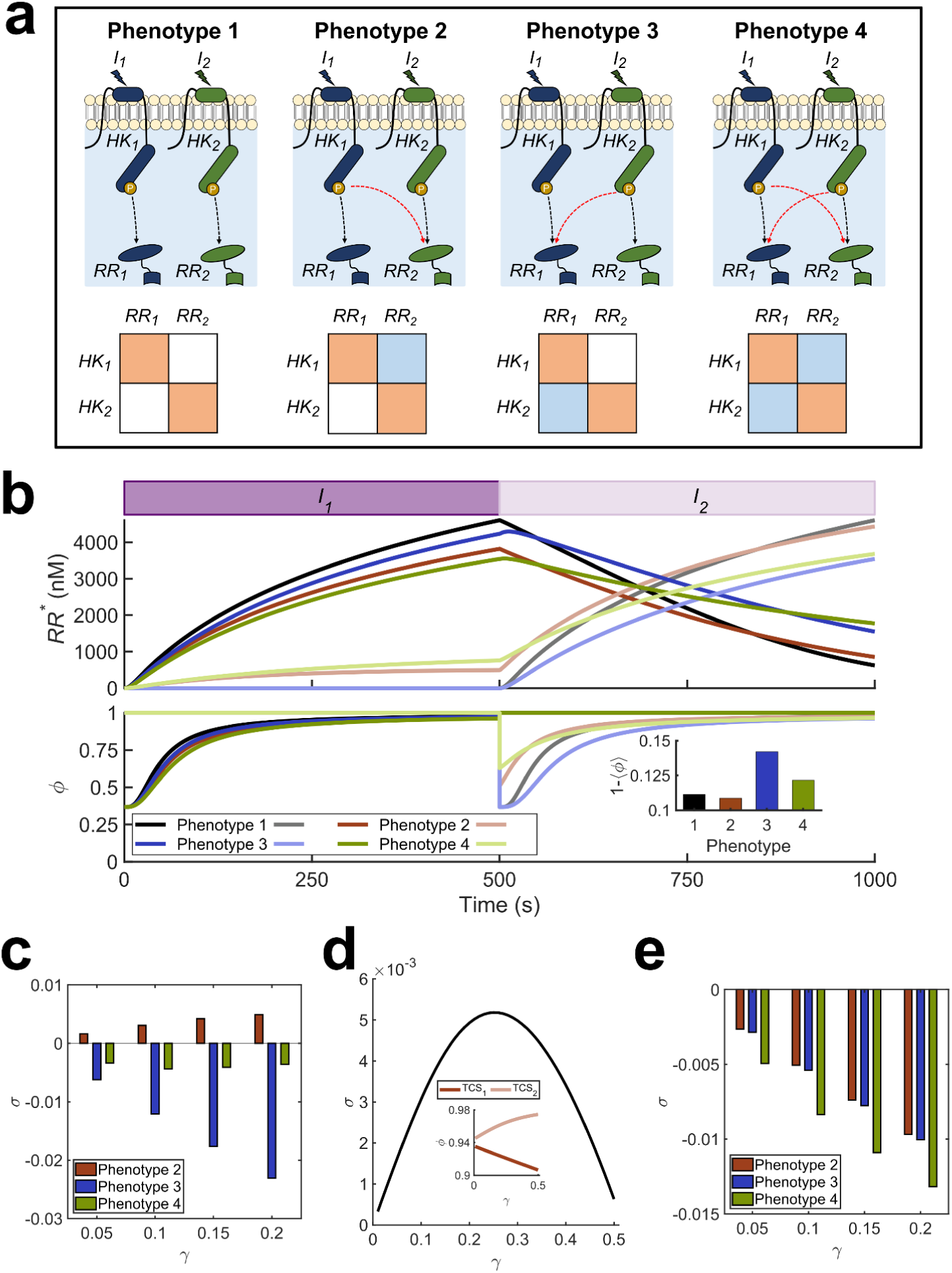
Mathematical model of TCS signaling predicts advantages of crosstalk. (**a**) All possible phenotypes with N=2 TCSs. Cognate interactions (black arrows) and crosstalk (red arrows) are shown. These interactions are also depicted compactly in the ‘interaction matrix’ for each phenotype (Methods). Orange squares represent cognate interactions and blue squares crosstalk. (**b**) Signal-response behavior and fitness of the phenotypes in a programmed environment. The purple filled rectangles depict the presence of the input signals, with the darker shade representing I_1_ and the lighter shade I_2_. The signal strength is 10^4^ nM for both. The top panel shows the concentrations of activated RRs and the bottom panel the associated fitness of the responses. The phenotypes are color coded and dark and light curves represent TCS_1_ and TCS_2_, respectively. Crosstalk strength is γ=0.26. The inset shows the reduction in time-averaged fitness of the different phenotypes due to the signals. The fitness is 1 in an unperturbed environment. (**c**) Selection coefficient in a programmed environment. σ as a function of γ when I_1_ is followed by I_2_. (**d**) Optimal crosstalk strength. Dependence of σ on γ for phenotype 2. Inset shows the fitness of the two TCSs contributing to σ. (**e**) Selection coefficients in random environment. σ as a function of γ when I_1_ and I_2_ follow no order. Fitness is calculated as the mean over all possible signal sequences.

For phenotype 1, where TCSs are insulated, our model predicted that the responses to the two signals were, expectedly, identical except for a shift in time (black curves in Fig. 1b). When I_1_ arrived, bacterial fitness dropped sharply, indicating a changed environment to which the bacterium was yet to adapt. The bacterium mounted an adaptive response, improving its fitness with time. As RR_1_-P increased, the fitness, *ϕ*_1_, recovered. The same phenomenon was observed upon the arrival of I_2_. The absence of crosstalk implied that the responses to I_1_ and I_2_ were independent. Although the fitness was nearly fully restored eventually, the time-averaged overall fitness, ⟨*ϕ*⟩, was lower than unity, indicative of the vulnerability of the bacterium ‘during’ adaptation to the changed environment.

For phenotype 2, with HK_1_→ RR_2_ crosstalk (red curves in Fig. 1b), our model predicted that before the arrival of I_2_, signal leakage to TCS_2_ resulted in lower RR_1_-P and, hence, *ϕ*_1_ than for phenotype 1. The signal leakage, however, triggered TCS_2_. The resulting RR_2_-P upregulated HK_2_ and RR_2_. When I_2_ came up, the bacterium responded faster and better than phenotype 1; RR_2_-P and *ϕ*_2_ were higher than for phenotype 1. The overall fitness, ⟨*ϕ*⟩, increased beyond that of phenotype 1. Thus, the bacterium was predicted to be more sensitive and responsive to the upcoming stimulus due to crosstalk, increasing its fitness. This scenario was illustrative of the possible advantage of crosstalk in a programmed environment.

For phenotype 3, with HK_2_→ RR_1_ crosstalk, in our model predictions, the needless signal dissipation to RR_1_ following the onset of I_2_ induced a fitness loss (blue curves in Fig. 1b). Finally, for phenotype 4, with bidirectional crosstalk, RR_1_-P was like phenotype 2 due to dissipation before the arrival of I_2_, but the advantage of priming was lost due to the HK_2_→ RR_1_ crosstalk after the arrival of I_2_, resulting in an overall fitness loss (green curves in Fig. 1b). The predicted time-averaged fitness loss, 1− ⟨*ϕ*⟩, of the four phenotypes over the entire signal-response period highlights the advantage of phenotype 2, which has a crosstalk pattern that mirrors the signal sequence, over the other phenotypes (Fig. 1b inset).

Next, we examined how the fitness advantage would depend on the strength of crosstalk using our model. We defined the selection coefficient, *σ*, for any phenotype as the difference between the time-averaged fitness of the phenotype and that of phenotype 1, the latter without any crosstalk. We quantified the strength of crosstalk using γ, the ratio of the rates of phosphotransfer to noncognate and cognate RRs (Methods). The larger was the value of γ, the greater was the extent of crosstalk. We found from our predictions that for all the values of γ studied, phenotype 2 had positive *σ*, whereas the other phenotypes had negative *σ* (Fig. 1c), consistent with the results above. Further, for phenotype 2, *σ* displayed a maximum at intermediate γ (Fig. 1d). Increasing γ increased priming and improved the response to I_2_, increasing fitness. Beyond a point, however, the advantage of priming diminished, but the response to I_1_ continued to be compromised, lowering the overall fitness (Fig. 1d inset). Thus, according to our model, limited crosstalk offered a fitness advantage to phenotype 2.

### Specificity is advantageous in ‘random’ environments

Using the same phenotypes above, we applied out model to estimate *σ* in a random environment, where there was no defined sequence of signals (Methods). Now, phenotype 1 had the highest estimated fitness; *σ* was negative for all the other phenotypes (Fig. 1e). Because the upcoming signal was not pre-specified, priming conferred no advantage. The detrimental effects of crosstalk then decreased fitness regardless of the crosstalk pattern. Thus, *σ* was equal for phenotypes 2 and 3, which had one crosstalk interaction each, and lower for phenotype 4, which had two crosstalk interactions. Moreover, the greater the value of γ, the lower was the value of *σ* in the random environment. Thus, in the absence of a consistent sequence of stimuli, our model predicted that evolutionary pressure may select for specificity.

Using sensitivity analysis, we found that the inferences above were robust to variations in parameter values (Supplementary Fig. 1). Furthermore, our findings were robust to the fitness construct employed (Supplementary Text 1; Supplementary Fig. 2) and the nature of the signals; we tested both square pulses and exponentially decaying signals (Supplementary Fig. 3). Our model also predicted that with decaying signals the fitness advantage of crosstalk ceased when the interval between the signals was either too small or too large (Supplementary Fig. 3). When the interval was too small, the second signal appeared before significant priming could happen, whereas when the interval was too large, the priming faded away before the second signal could arrive. These latter predictions were consistent with observations in *E. coli* (23), where priming conferred a significant fitness advantage, manifested as enhanced growth rate, only for a range of time gaps between signals.

**Fig. 2.**
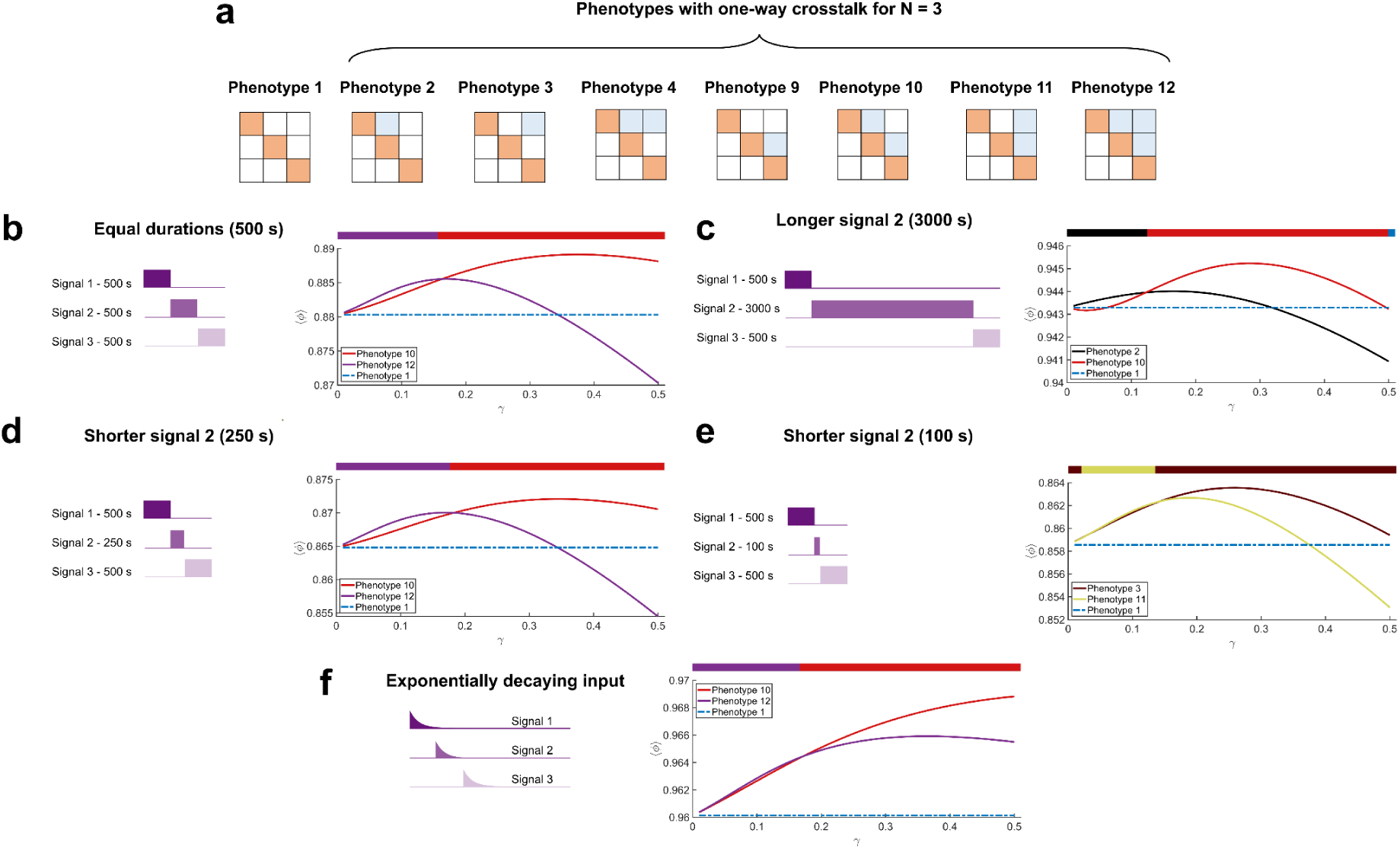
One-way crosstalk patterns yielded the fittest phenotypes. (**a**) One-way crosstalk patterns with N=3 TCSs. Interaction matrices of phenotype 1, without crosstalk, and seven other phenotypes with different one-way crosstalk patterns. The signal sequence is I_1_→ I_2_→ I_3_. The fitness of the fittest phenotypes and of phenotype 1 as functions of the strength of crosstalk, γ, when (**b**) signals were of the same duration (500 s), or when I_2_ lasted (**c**) 3000 s, (**d**) 250 s, and (**e**) 100 s, and (**f**) when the signals decayed exponentially. The colored bars at the top of each panel graphically depict the range of γ over which the respective color-coded phenotype has the highest fitness. Cartoons of the signal patterns are at the left in each panel.

**Fig. 3.**
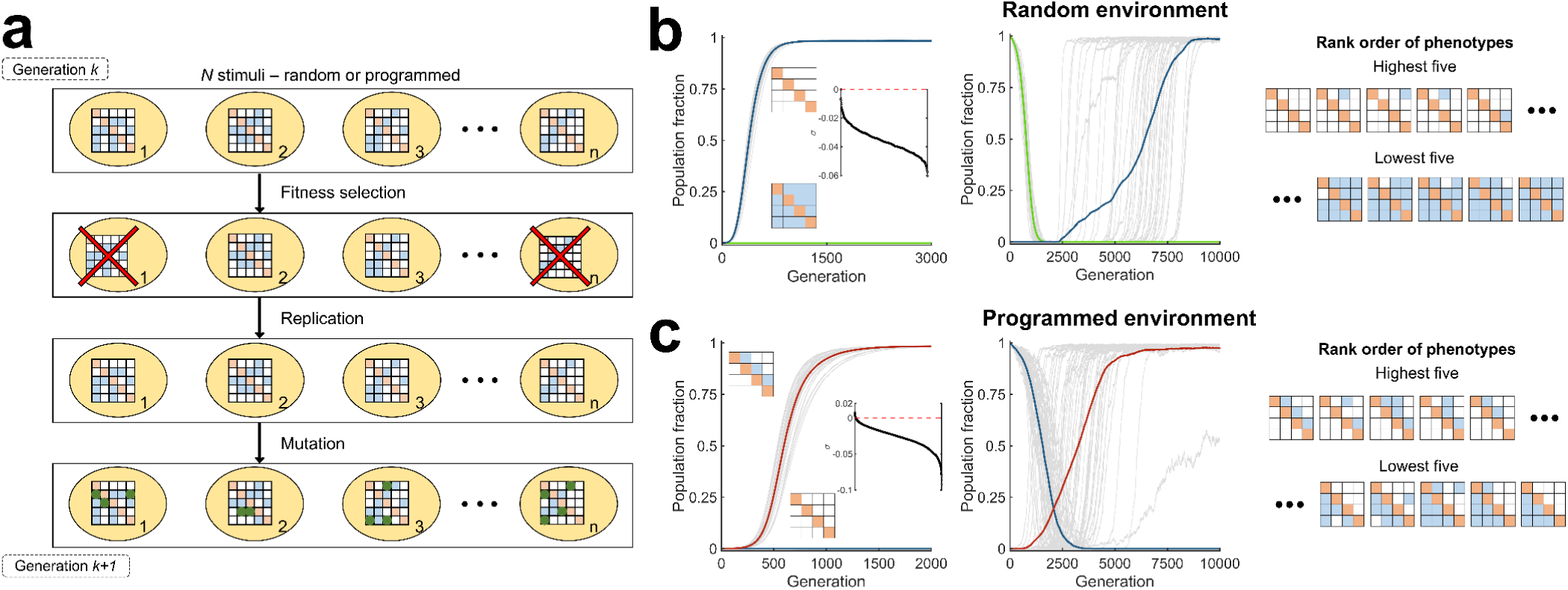
Stochastic evolutionary dynamics simulations show selection of crosstalk in programmed environments and specificity in random environments. (**a**) Schematic of Wright-Fisher simulations. Simulations proceed in discrete generations and with fixed populations (n) comprising bacteria of different phenotypes, indicated by their interaction matrices. In each generation, bacteria are exposed to stimuli. Depending on their response, fitness selection takes place and less-fit bacteria are eliminated. Lost bacteria are replaced with copies of selected ones, chosen randomly. The resulting bacteria mutate, illustrated using green boxes in the interaction matrices, resulting in altered phenotypes, which form the substrate for selection in the next generation. (**b**) Evolution in a random environment. The phenotype without any crosstalk (blue) gets fixed whether the initial population is homogeneous (left) or mixed (middle). The phenotype with all crosstalk interactions is also shown for comparison (green). The gray lines are trajectories of the two phenotypes in each of 50 realizations. The thick lines are means. Trajectories of all other phenotypes are not shown. The crosstalk strength was set to γ = 0.26. The inset in the left plot is the rank-ordered selection coefficient of all the phenotypes. The interaction matrices of the five most and five least fit phenotypes are shown (right). (**c**) Evolution in a programmed environment. The one-way crosstalk phenotype mirroring the signal sequence I_1_→ I_2_→ I_3_→ I_4_, which has the highest fitness, dominates the population (red), whether the initial population is homogeneous (left) or mixed (middle). The inset in the left plot is the rank-ordered selection coefficient of all the phenotypes. The interaction matrices of the five most and five least fit phenotypes are depicted (right). Simulations used N=4 TCSs.

### Programmed environments favor one-way crosstalk

For the minimal case of N=2, phenotype 2 alone could anticipate I_2_ following I_1_ and thus was predicted to have the highest fitness in our model. For bacteria with more than two TCSs, the fittest phenotype is not obvious, as such anticipation is possible with multiple phenotypes. For instance, the phenotype with the crosstalk interactions HK_1_→ RR_2_ and HK_2_→ RR_3_ as well as the phenotype with HK_1_→ RR_2_ and HK_1_→ RR_3_ could anticipate the sequence I_1_→ I_2_→ I_3_. The number of phenotypes grows exponentially with N. A bacterium with N TCSs will have N cognate and up to N(N-1) noncognate interactions. Depending on whether each of the latter interactions is realized or not, a total of 2^N(N-1)^ phenotypes can exist, each representing a distinct crosstalk pattern. For N=3, this would amount to 2^6^=64 phenotypes and for N=4 to 2^12^=4096 phenotypes. Identifying the fittest phenotype would thus require a comprehensive assessment of each of these phenotypes. We performed this assessment next.

We considered N=3. We numbered the phenotypes from 1 to 64, starting with the phenotype with no crosstalk and ending with the phenotype with all crosstalk interactions realized (Fig. 2a). We subjected each phenotype to a programmed environment with the signal sequence I_1_→ I_2_→ I_3_. We also allowed the signals to have different durations, more realistically mimicking natural environments. For each scenario, we applied our model to predict signal-response characteristics and estimated the resulting fitness.

When the signals were all of the same duration, our model predicted that the phenotype that was the fittest depended on the strength of crosstalk, γ. When γ was small, phenotype 12, which had HK_1_→ RR_2_, HK_2_→ RR_3_ and HK_1_→ RR_3_ interactions was the fittest (Fig. 2b). Its fitness was only slightly higher than that of phenotype 10, which had HK_1_→ RR_2_ and HK_2_→ RR_3_ interactions. Note that both these phenotypes anticipated upcoming signals and were fitter than phenotype 1, which had no crosstalk. As γ increased, phenotype 10 became fitter than phenotype 12 in our predictions. Interestingly, the fitness of the latter decreased beyond a threshold γ and eventually dropped below that of phenotype 1. Phenotype 10, however, remained fitter than phenotype 1 throughout. We understood these trends as follows. When γ was low, the cost of signal dissipation was small. Thus, the gain from crosstalk by HK_1_ with both RR_2_ and RR_3_ and by HK_2_ with RR_3_ more than compensated for the fitness loss due to leakage. However, as γ increased, the latter cost increased and limiting crosstalk became advantageous. Accordingly, our model predicted that crosstalk between HK_1_ and RR_2_ and between HK_2_ and RR_3_, which ensured the requisite anticipation of upcoming signals, were retained, resulting in an overall fitness gain, whereas the redundant crosstalk between HK_1_ and RR_3_ was eliminated in the fittest phenotype.

We next increased the duration of I_2_ 6-fold (Fig. 2c). When γ was small, phenotype 2, which had the HK_1_→ RR_2_ interaction alone was the fittest in our predictions. As γ increased, phenotype 10, which had HK_1_→ RR_2_ and HK_2_→ RR_3_ interactions, became the fittest. With weak crosstalk, the advantage of priming to I_3_ through the entire duration of I_2_ was not enough to compensate for the loss of response to I_2_. Phenotype 2, which did not have the HK_2_→ RR_3_ interaction was therefore the fittest. On the other hand, when crosstalk was stronger, the priming from both HK_1_→ RR_2_ and HK_2_→ RR_3_ compensated for any signal dissipation, rendering phenotype 10 the fittest in our predictions.

We also considered the effect of shortening the duration of I_2_ (Fig. 2d, e). When the duration was shortened by 50%, phenotypes 12 and 10 were predicted to be the fittest, depending on γ, in a manner similar to when the signals were all of the same duration (Fig. 2b, d). The shortening of the duration by 50% thus did not affect the cost-benefit analysis substantially. Shortening the duration 5-fold, however, made a difference, with phenotypes 3 and 11 now the fittest (Fig. 2e). As above, when γ was small, phenotype 11, with the crosstalk interactions HK_1_→ RR_3_ and HK_2_→ RR_3_, both anticipating the upcoming signal I_3_, was the fittest in our model. This was because at low values of γ, priming to I_3_ while I_2_ was present did not add to the cost due to signal dissipation significantly, as I_2_ was present for a short while. However, as γ increased, phenotype 3, which had the single crosstalk interaction HK_1_→ RR_3_ was the fittest. The cost of dissipation, although I_2_ was short-lived, was no longer affordable. The phenotype that let I_1_ prime the bacterium to the next ‘major’ signal, I_3_, was thus the fittest. Finally, as with the N=2 scenario, the results were similar when exponentially decaying signals were used instead of square pulses (Fig. 2f).

In all these cases, an intriguing feature of the fittest phenotypes is directed, ‘one-way’ crosstalk. If we denote the signal sequence as I_1_→ I_2_→ I_3_→ …, then the fittest phenotypes had crosstalk of the type HK_i_→ RR_j_ with j>i. In other words, the crosstalk that enabled priming to ‘upcoming’ signals was favored. Reverse signal flow, where j<i, resulted in phenotypes that suffered fitness loss. In the interaction matrices, the fittest phenotypes all had non-zero entries in the upper triangular portions and never in the lower triangular portions (Fig. 2a). To test the robustness of this prediction, we adopted two strategies. We performed comprehensive evolutionary dynamics simulations to examine whether the fitness advantage predicted by the calculations above would lead to the selection of the corresponding phenotypes with the one-way crosstalk patterns. Second, we sought evidence of these predictions in available experimental data.

### Evolutionary simulations predict selection of phenotypes with one-way crosstalk patterns mirroring signal sequences

Using the descriptions above of the responses of different phenotypes to stimuli, we performed stochastic, discrete generation, Wright-Fisher evolutionary simulations (24) (Fig. 3a; Methods) to determine which phenotypes would get selected in different environments. We now considered N=4 TCSs, increasing the complexity to a total of 4096 phenotypes, making it even more difficult to predict the fittest phenotypes intuitively. We performed simulations with two types of initial conditions: 1) the ‘homogeneous condition’, where a single phenotype existed, and 2) the ‘mixed condition’, where all the phenotypes were equally represented. With each initial condition, we considered both random and programmed environments. With N=4, we had four types of signals, one for each of the TCSs. We let each bacterium be stimulated four times. In the random environment, each stimulus was chosen randomly from the four possible signals. In the programmed environment, the signals followed a predetermined sequence, where the signals all appeared once and in a fixed order. We computed the fitness of each of the 4096 species in each of these environments. In each generation, we allowed every bacterium to be selected with a probability proportional to its fitness. The selected bacteria were duplicated to replace lost bacteria and ensure a constant bacterial population. The bacteria were then subjected to mutations. A mutation involved a change in the crosstalk network of the bacterium, resulting in an altered phenotype. Specifically, we allowed each of the N(N-1)=12 potential crosstalk interactions within a bacterium to be flipped (from existent to non-existent and *vice versa*) with a probability μ, the mutation rate, in each generation. The resulting pool of bacteria formed the substrate for evolution in the next generation. We repeated this process over a large number of generations and performed several realizations.

In the random environment, our simulations predicted that the phenotype without any crosstalk dominated the population (Fig. 3b). For the homogenous condition, we initiated simulations with the species containing all crosstalk interactions. Gradually, phenotypes with fewer crosstalk interactions emerged. Eventually, the phenotype with no crosstalk emerged and dominated the population. With the mixed condition, the latter species began to dominate the population from the early stages and was soon fixed in the population. These observations agree with the prevalent paradigm of TCS signaling favoring specificity (5, 8, 9, 12). Also, rank-ordering phenotypes by their fitness values (Fig. 3b, inset) revealed that phenotypes with increasing number of crosstalk interactions had decreasing fitness. To illustrate this, we present the crosstalk patterns of the top five and bottom five fittest phenotypes (Fig. 3b). The former have zero or one crosstalk interaction and the latter have all or one less crosstalk interactions, respectively.

In the programmed environment, which followed the signal sequence I_1_→ I_2_→ I_3_→ I_4_, the phenotype with the crosstalk pattern mirroring this signal sequence dominated the population (Fig. 3c). For the homogeneous condition, we used the species without crosstalk to initiate simulations. Gradually, mutants with crosstalk emerged and grew, causing the initial species to decline. Eventually, the phenotype with the crosstalk pattern mirroring the signal sequence emerged and dominated the population. For the mixed condition, the latter phenotype grew from the early stages and was rapidly fixed. Arranging the fitness values in descending order (Fig. 3c, inset) displays the benefit of priming for upcoming stimuli. The five fittest phenotypes all had crosstalk interactions in the upper triangle of their interaction matrices, indicating one-way crosstalk patterns that prime bacteria to upcoming signals (Fig. 3c). The least fit phenotypes had the lower triangle of the interaction matrices populated, indicating crosstalk that had signal flows opposite to the sequence of stimuli.

These results were not restricted to N=4 TCSs. With N=2 (Supplementary Fig. 4) and N=3 TCSs (Supplementary Fig. 5) as well, the phenotype with no crosstalk was selected in random environments and the phenotype with the crosstalk pattern mirroring the sequence of signals was selected in programmed environments.

**Fig. 4.**
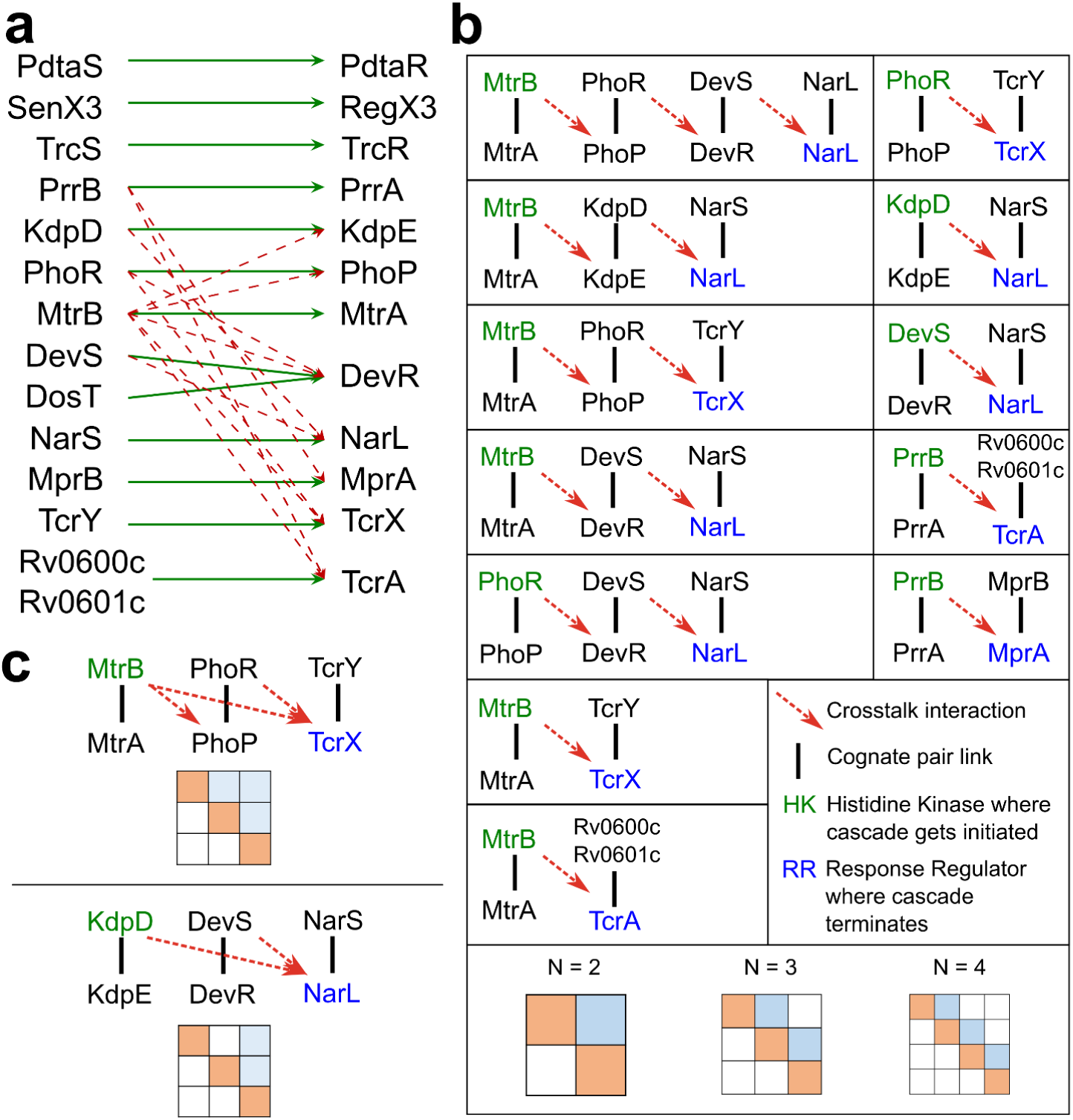
Crosstalk patterns in *M. tuberculosis* TCSs *in vitro* were one-way. (**a**) Complete crosstalk map between TCSs of *M. tuberculosis*. The HKs (left column) and their cognate RRs (right column) are connected by green arrows. Crosstalk interactions observed (15) are shown as red dashed arrows. (**b**) Crosstalk cascades. All possible signal flows based on the crosstalk interactions in (**a**). (**c**) Superimposed signal cascades. Examples of crosstalk patterns resulting from superimposition of cascades from (**b**).

**Fig. 5.**
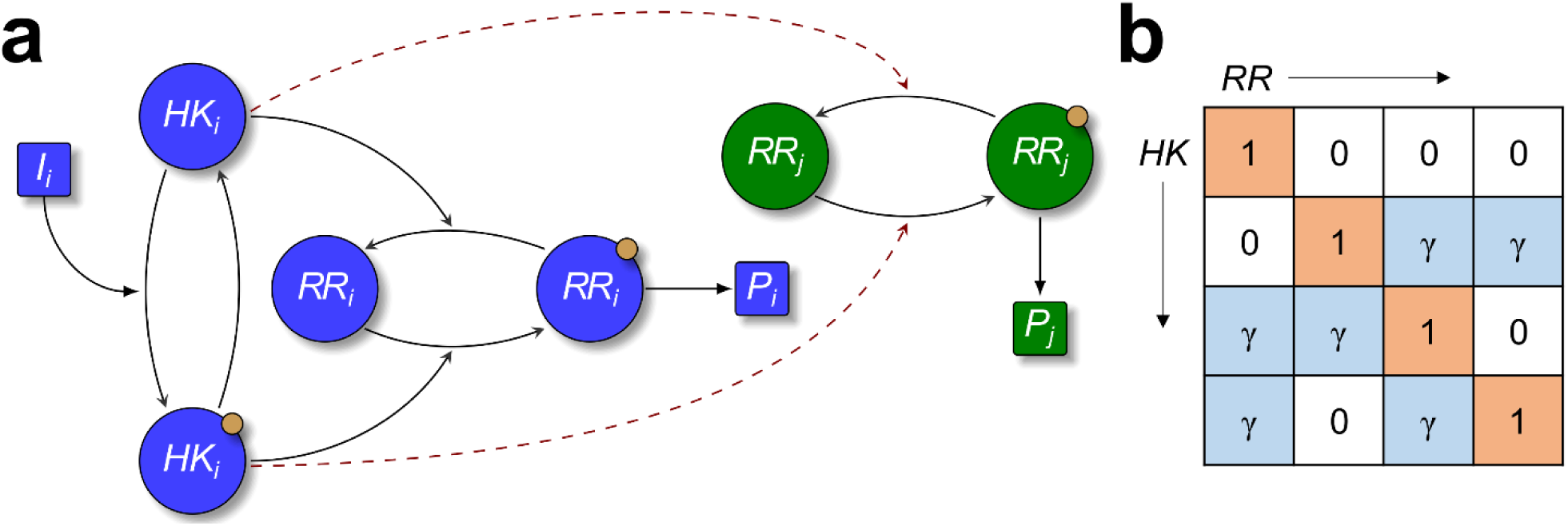
Schematic of mathematical model of TCS signaling with crosstalk. (**a**) Architecture of the generalized mathematical model. The input I_i_ is detected by HK_i_, which gets phosphorylated (HK_i_ with a yellow dot) and then transfers the phosphoryl group either to the cognate response regulator, RR_i_ (blue), or non-cognate response regulator (RR_j_, *j ≠ i* (green)). Activated RRs trigger downstream gene expression via promoter P_i_. Inactive HKs can act as phosphatases, which dephosphorylate active RRs. (**b**) Sample interaction matrix for N=4. The diagonal positions represent cognate and non-diagonal positions non-cognate interactions. Zeros in the non-diagonal cells represent the absence of the corresponding crosstalk interactions. The ratio of the phosphotransfer rate for non-cognate and cognate interactions is γ. 2^N(N-1)^ such interaction matrices are possible depending on whether each non-diagonal entry is zero or not.

These simulations thus point to environments where crosstalk may be evolutionarily favored. It is possible that such programmed environments may have been the reasons for the selection of the crosstalk that is observed in some bacteria. Our model and simulations go beyond offering a plausible explanation of the origins of such crosstalk and predict that the crosstalk selected is expected to be one-way. We next sought evidence of one-way crosstalk patterns in available experimental data.

### Evidence of one-way crosstalk in TCSs of *M. tuberculosis*

In a recent study, crosstalk between the TCSs of *M. tuberculosis* has been mapped using *in vitro* assays of phosphotransfer from HKs to all cognate and non-cognate RRs (15). Significant crosstalk was observed (Fig. 4a), which allowed us to assess signal flows through extended TCS networks. Using the crosstalk interactions, we identified all possible signal flows, or cascades, in the TCSs of *M. tuberculosis* as follows. We considered the HK PhoR, for instance, which showed crosstalk with the RR DevR (Fig. 4a). DevS, the cognate HK of DevR, further showed crosstalk with the RR NarL. NarS, the cognate HK of NarL, did not engage in any crosstalk. Thus, when PhoR gets activated, it can transmit a portion of the signal to DevR. Similarly, crosstalk of DevS with NarL would transmit some portion of the signal from DevS-DevR to the NarS-NarL system, at which point the signal flow would be terminated. Hence, PhoR-PhoP, DevS-DevR, and NarS-NarL form a cascade of signal flow via crosstalk. In this cascade, the signal is not transmitted either to PhoP from DevS or NarS or to DevR from NarS, making the flow one-way.

Following the procedure above, we started with each of the TCSs of *M. tuberculosis* and traced the resulting cascades. We found 12 such cascades (Fig. 4b). The longest cascade involved 4 TCSs. There were four cascades involving 3 TCSs each and seven cascades involving 2 TCSs each. (Representative interaction matrices for all these cases are presented at the bottom in Fig. 4.) Note that all the cascades had one-way crosstalk with the patterns resembling the fittest phenotypes in our simulations above.

By superimposing the cascades above, we obtained additional one-way crosstalk patterns, reflective of the patterns identified in our simulations. Two such patterns are depicted in Fig. 4c. For instance, the crosstalk pattern involving MtrB-MtrA, PhoR-PhoP, and TcrY-TcrX (Fig. 4c top panel) was equivalent to phenotype 12 in the N=3 case discussed above (Fig. 2b). Similarly, the pattern involving KdpD-KdpE, DevS-DevR, and NarS-NarL (Fig. 4c bottom panel) was equivalent to phenotype 11 in the N=3 case above (Fig. 2b). Remarkably, we could not find any crosstalk pattern that was not one-way. This evidence of exclusive one-way crosstalk in the TCSs of *M. tuberculosis* offered strong support to the predictions of our model and simulations.

To assess whether the crosstalk could have evolutionarily underpinnings, we sought signatures of evolutionary pressures against diversification post gene duplication in the sequences of the TCS proteins using bioinformatics analysis (Supplementary Text 2). The analysis, conducted on a subset of the TCSs, suggested that this evolutionary pressure may have been lesser for the TCSs involved in crosstalk than for the TCSs that were specific, offering further support to the notion that the observed crosstalk may have been evolutionarily favored (Supplementary Text 2, Supplementary Fig. 6 and 7, Supplementary Table 2).

## DISCUSSION

Despite the strong evolutionary arguments favoring specificity in bacterial TCSs (4, 5), crosstalk between TCSs has been observed (14, 15). Here, we present an alternative evolutionary paradigm where crosstalk would be advantageous. Using modeling of TCS signaling networks and comprehensive evolutionary dynamics simulations, we predicted that in programmed environments, where stimuli arrive in a predetermined sequence, crosstalk that would prime bacteria to upcoming signals would confer an evolutionary benefit. Thus, specific crosstalk patterns that mirror the sequences of stimuli could get selected in bacteria living in such environments. Analyzing recent *in vitro* data (15), we found that potential crosstalk networks involving the TCSs of *M. tuberculosis* all displayed one-way signal flow, consistent with the notion of priming and selection in programmed environments. This new evolutionary paradigm is not in conflict with the paradigm underlying specificity. Our modeling and simulations predicted that when no predetermined sequence of stimuli existed, specificity was evolutionarily favored. Our study, thus, offers a conceptual framework that synthesizes specificity and crosstalk in bacterial TCS systems. They appear to be two sides of the same coin; they are both outcomes of the same evolutionary forces, but in environments that present signals differently. Programmed environments may be rarer, resulting in the lower prevalence of crosstalk.

Independent evidence exists of one-way crosstalk aiding bacterial adaptation in programmed environments. In *E. coli*, evolutionary experiments showed how ‘anticipation’, facilitated by crosstalk, is selected for when the environment displays a specified pattern of carbon source switching (22). Similarly, in *S. cerevisiae*, preparation of the bacterium to respond to oxidative stress while experiencing heat shock was a result of adaptation; these stresses are typically experienced in the same temporal order (22). Furthermore, the complex structure of environments can become ingrained in *in silico* biochemical networks in order to predict environmental changes preemptively (25). In agreement, this adaptive behavior was evident in *E. coli*, where a match between the covariation of transcriptional responses and the sequence of temperature and oxygen stresses triggering them was observed (25). Evidence also exists of pathogenic bacteria evolving crosstalk to adapt to their hosts. For instance, mutations in the TCS BfmS-BfmR of *P. aeruginosa* in individuals with cystic fibrosis were recently found to alter, facilitated via crosstalk by the noncognate HK GtrS, regulation of downstream gene expression in order to promote biofilm formation and chronic infection (26). Similarly, in α-proteobacteria, multiple HKs of the HWE/HisKA-2 family can control the phosphorylation of the same response regulators in a coordinated manner and tune downstream gene expression (27).

Based on the signaling cascades we deduced from the *in vitro* TCS crosstalk interactions of *M. tuberculosis*, it would be interesting to identify corresponding sequences of stimuli, potentially unveiling information of the environments to which *M. tuberculosis* may have adapted. The ligands/stimuli that many of the TCSs sense, however, remain unknown, precluding such analysis (28). Yet, specific instances suggesting such adaption could be identified from the cascades. For example, the TCS PrrB-PrrA is reported to be involved in the early replication steps of *M. tuberculosis* inside macrophages (29). The TCS MprB-MprA has been argued to be essential for establishing persistent infection (30), a state of slower or halted replication from which the bacterium can be reactivated to establish active infection (31). Disruption of *mprA* affected processes required for survival during the persistence and subsequent infection stages (30). One could thus argue that crosstalk from PrrB-PrrA to MprB-MprA may be favorable because it would prime the bacterium to activate the processes necessary for establishing persistent infection, a key feature of successful tuberculosis infection (32), once entry is gained into a macrophage. Indeed, this one-way crosstalk was observed in the *in vitro* cascades (15). Future experiments may assess the advantage of such crosstalk *in vivo*.

Crosstalk is not limited to bacterial TCSs. Examples of crosstalk exist in human growth factor signaling networks (33), MAPK networks of yeast (34), and between TOR and CIP pathways in *S. pombe* (35). The evolutionary underpinnings of these crosstalk interactions may be more difficult to unravel because of the more involved regulatory structures in these organisms than in the simpler bacterial TCS systems. Yet, controlled evolutionary experiments suggest selection of cross-regulation patterns in broad agreement with our predictions. For instance, the yeast *S. cerevisiae*, which is commonly used in the fermentation industry, is subjected to heat, ethanolic stress and oxidative stress, in that order, in the industrial process (22). The related regulatory networks were observed to have the following crosstalk interactions: heat→ ethanolic, heat→ oxidative, and ethanolic→ oxidative (22). This is similar to the phenotype 12 in the N=3 case in our model (Fig. 2a). Furthermore, when the organism was artificially exposed to these stresses in the reverse order, the crosstalk interactions switched their directions (22). These scenarios, together with our proposed paradigm, point to the possible evolutionary advantages of crosstalk.

Because of its evolutionary advantages, crosstalk may be a potential target of intervention. With pathogenic bacteria, crosstalk may sharpen the already sophisticated strategies to evade host immune responses and promote virulence (28, 36). Bacterial HKs offer promising targets of intervention (1, 18). Where crosstalk may aid bacterial survival and adaptation, as suggested for instance with *M. tuberculosis* (15), targeting HKs engaged in crosstalk could prove a more potent strategy than targeting specific HKs. It would not only block the cognate response of the targeted HK, but also compromise the responses of the TCSs that would otherwise have been primed by the targeted HK via crosstalk.

## METHODS

### Mathematical model of TCS signaling with crosstalk

We developed a mathematical model to describe bacterial signal transduction via TCSs. We considered the scenario in which a bacterium contains N distinct TCSs, which can be engaged in crosstalk (Fig. 5a). We built the model by envisioning the set of events associated with the *i*^th^ TCS engaged in crosstalk with the *j*^th^ TCS (*i, j* ∈{1, 2,…, *N*}), listed below as reactions.

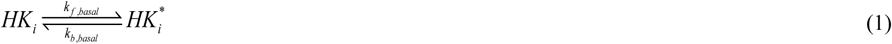

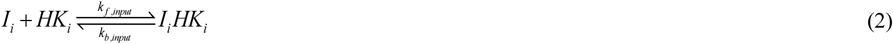

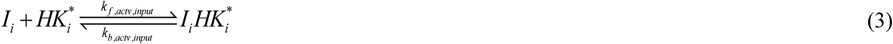

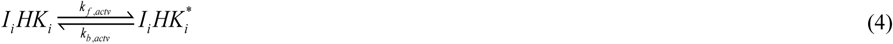

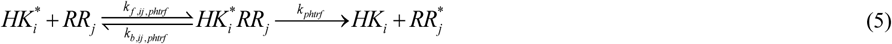

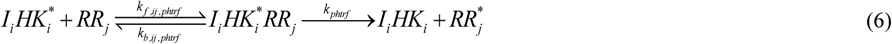

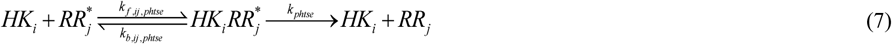

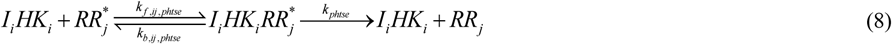

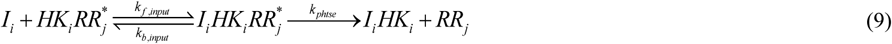

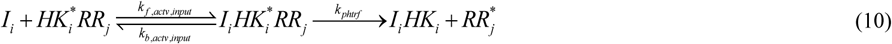

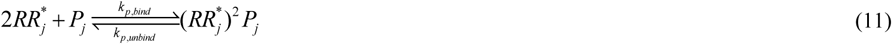

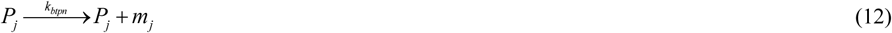

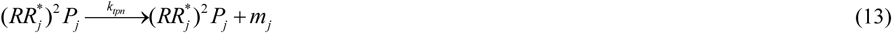

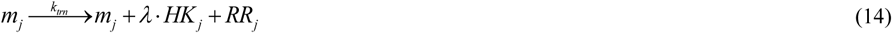

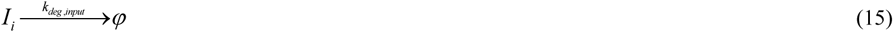

Here, the subscript *i* refers to the *i*^th^ TCS. We recognize that *HK*_*i*_ can be activated reversibly at some basal level, *i*.*e*., in the absence of any input signal, to its active form,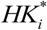 (reaction (1)) (37). The input, *I*_*i*_, can bind reversibly to *HK*_*i*_ or 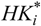 to yield the complexes *I*_*i*_*HK*_*i*_ or 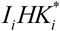, respectively (reactions (2) and (3)). *I*_*i*_*HK*_*i*_ can lead to the activated complex 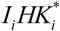 at a rate higher than the basal rate above (reaction (4)). 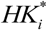 can bind *RR*_*j*_ and activate it via phosphotransfer, yielding *HK*_*i*_ and 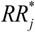 (reaction (5)). An analogous reaction occurs with 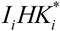 binding to *RR*_*j*_ (reaction (6)). Note that in these reactions, *j*=*i* would imply cognate interactions. *HK*_*i*_ can bind to 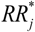 and exert phosphatase activity (reaction (7)), consistent with the bifunctional nature of typical HKs, which act as both kinases and phosphatases (1, 9, 38). The latter activity can also be triggered by *I*_*i*_*HK*_*i*_ (reaction (8)). The reversible binding of *I*_*i*_ to the intermediate *HK-RR* complexes is also possible (reactions (9) and (10)). Thus, we assumed that *RR* binding to *HK* does not influence ligand binding to *HK*. Binding rates of non-cognate partners (*k*_*f,ij,phtrf*_ and *k*_*f,ij,phtse*_) are weaker than cognate partners (*k*_*f,ij,phtrf*_ and *k*_*f,ij,phtse*_), and the attenuation factor is 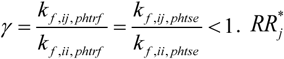 dimerizes and binds to the corresponding promoter *P*_*j*_ (reaction (11)). This binding enhances transcription compared to its basal level (reactions (12) and (13)); i.e., *k*_*tpn*_ *> k*_*btpn*_. Transcription produces mRNA, denoted by *m*, which are then translated, with the HK and RR translated in the ratio *λ* :1 (reaction (14)). Here, we recognize that the response also typically upregulates the corresponding TCS proteins (2, 39). Input signals degrade with rate constant *k*_*deg,input*_ (reaction (15)). All the other entities present in the network are assumed to degrade with a rate constant *k*_*deg*_ (not written explicitly for convenience).

Next, we estimated the rate of synthesis of *HK* and *RR* proteins by assuming that the DNA binding reactions are fast compared to transcription and translation reactions (15, 20). Let *P*_*T*_ be the total concentration of promoter binding sites present on the bacterial genome, with *f*_*b*_ and *f*_*f*_ the fractions of promoter sites in the bound and the free states, respectively. We assumed pseudo-equilibrium between DNA binding reactions, yielding

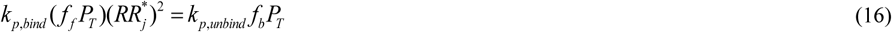

If *K*_1_ *= k*_*p,unbind*_ / *k*_*p,bind*_ is the equilibrium dissociation constant for reaction (11), we get

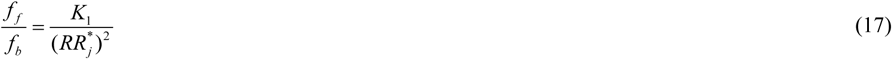

Because *f*_*b*_ + *f*_*f*_*=*1, it follows that

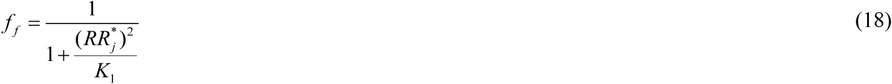

and

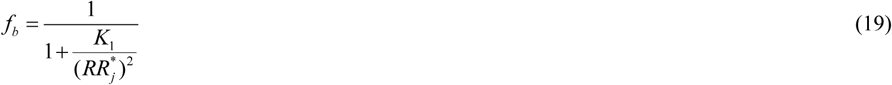

We now have the concentration of promoters in the basal and active states. Reactions (11) to (13) estimate the rate of upregulation of the corresponding TCS as follows. From reactions (12) and (13), the change of mRNA concentration would be

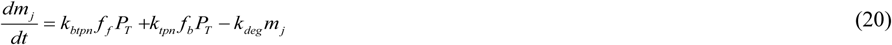

Applying the pseudo-equilibrium approximation to mRNA dynamics, *i*.*e*., 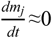, gives

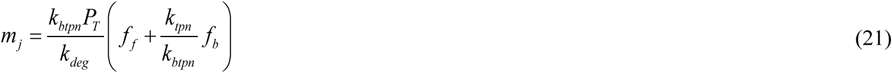

By substituting expressions (18) and (19) into (21), we obtain

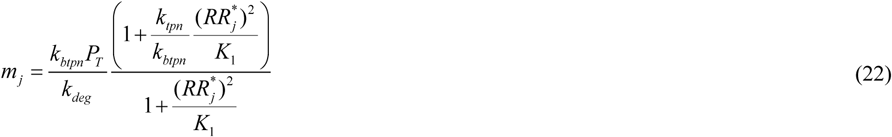

These mRNA molecules translate at the rate *k*_*trn*_ to produce *HK*_*j*_ and *RR*_*j*_ molecules in the ratio *λ*:1.

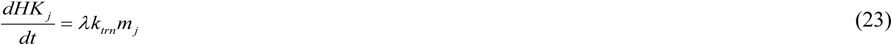

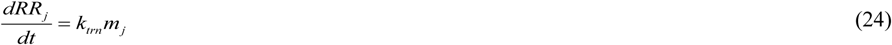

Substituting 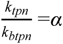 and 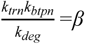, we get the synthesis rates of *HK* and *RR* by mRNA translation as

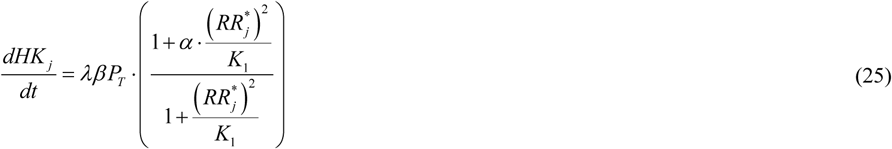

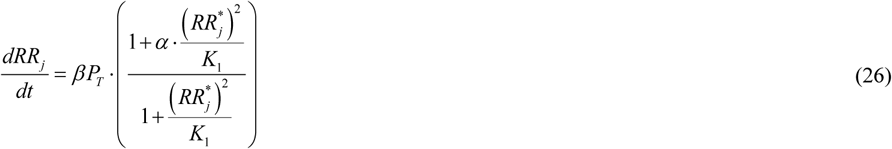

The rate equations for reactions (1) – (15) can be written following standard mass action terms and by utilizing the expressions (25) and (26) as follows.

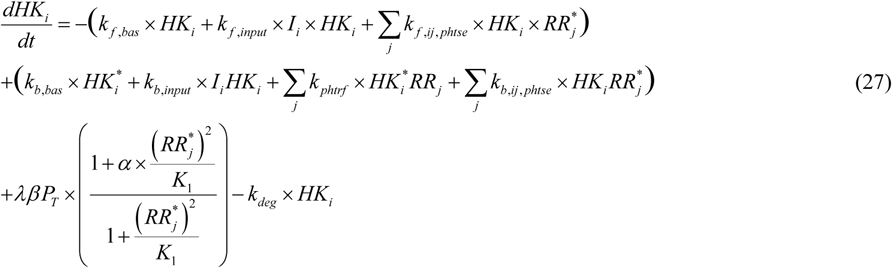

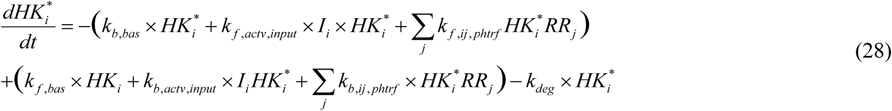

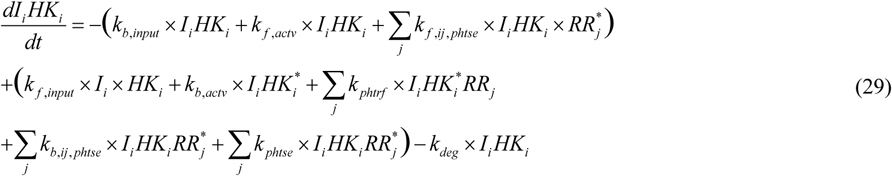

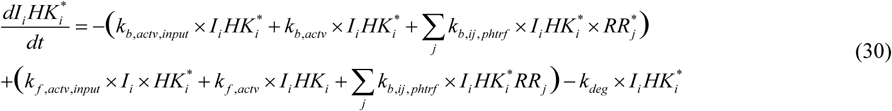

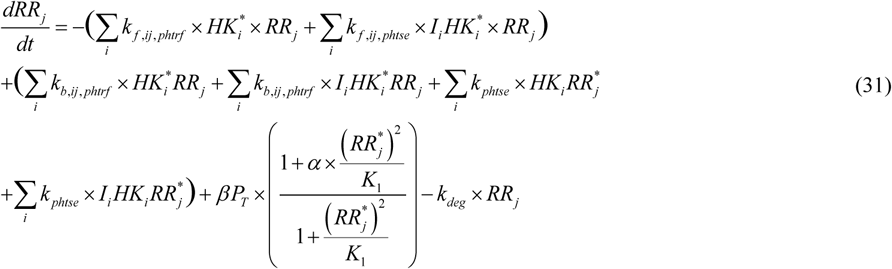

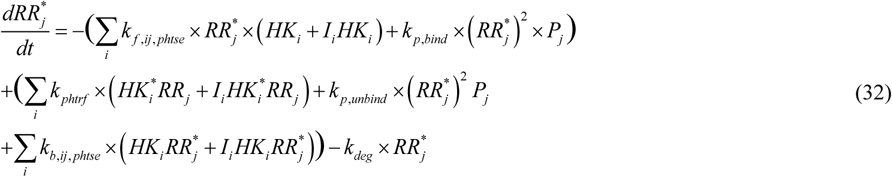

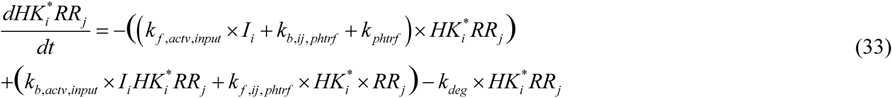

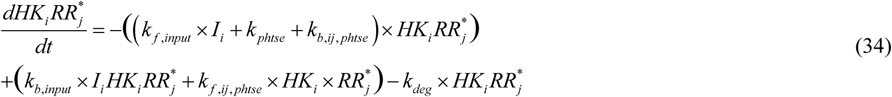

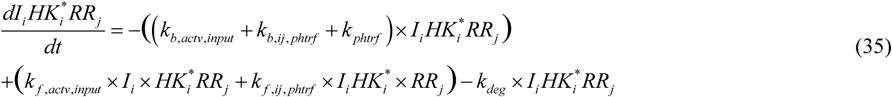

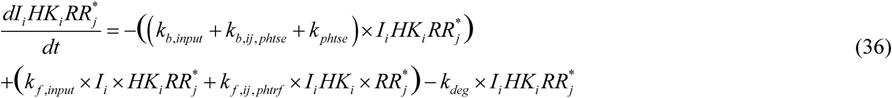

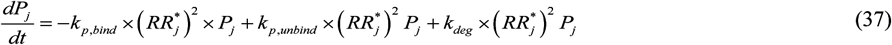

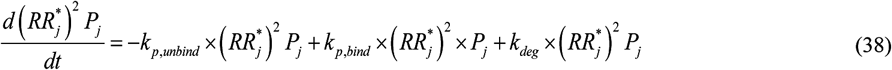

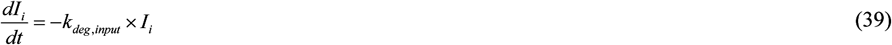

The rate constants involved were obtained from the literature (9, 20, 40) (Supplementary Table 1). The rate equations were integrated in MATLAB using the routine ode15s and with chosen initial conditions (Supplementary Table 1). In all our simulations, the above equations were first solved in the absence of stimuli for a sufficiently long time so that the basal autophosphorylation reactions balanced the degradation reactions and all the proteins reached a steady state. Using the latter as the pre-stimulus state of the bacterium, the above equations were solved in the presence of stimuli. The solution depended on the phenotype, described next.

### Interaction matrix

For a bacterium with N TCSs, different phenotypes are possible depending on the presence or absence of specific crosstalk interactions. An interaction matrix defines the identity of each phenotype (Fig. 5b). The *ij*^th^ element in the matrix represents the strength of the cross-interaction between *HK*_*i*_ and *RR*_*j*_ relative to the cognate interaction. The cognate interactions are all assumed to be equally strong and occupy the diagonal entries. The cross-interactions are also assumed to be of the same relative intensity, γ, whenever they exist. The non-diagonal entities are thus either 0 or γ. Since there are N(N-1) non-diagonal elements present, with 2 state values possible for each of them, we get 2^N(N-1)^ different phenotypes.

### Fitness formulation

We constructed a fitness variable based on the response of a TCS to a time-dependent input. We defined the fitness corresponding to the *i*^th^ TCS as

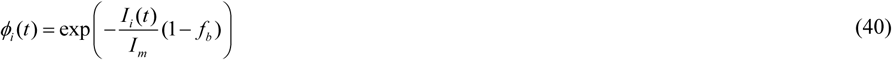

Where 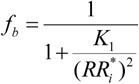 follows from Eq. (19) above. The term − *I*_*i*_ (*t*)/*I*_*m*_ reflects the inverse relationship between the fitness and input intensity. *I*_*m*_ is taken as the maximum (or peak) input value. Thus, as *I*_*i*_ increases, it reflects an increasing change in the environment, inducing a more significant fitness loss until the bacterium responds and adapts. The recovery of fitness following the response is determined by the second entity in the fitness variable, 1 − *f*_*b*_, where *f*_*b*_ denotes the fraction of promoters bound by *RR*^*\**^. (We recall that *K*_1_ is the dissociation constant of 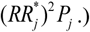 As this fraction increases, the magnitude of the response also rises, leading to greater fitness given the signal. This formulation of fitness makes sure that *ϕ*_*i*_ lies between 0 and 1. TCSs are assumed to contribute independently to fitness. Thus, for a bacterium with N TCSs, the total instantaneous fitness is the product of individual fitness values:

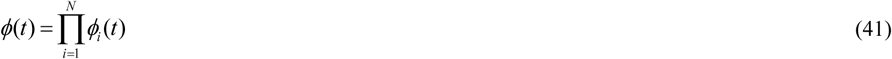

In the absence of any signal, *ϕ =* 1. Similarly, with a perfect response, *i*.*e*., with *f*_*b*_ *=* 1, *ϕ* is again 1. We also considered an alternative fitness formulation and found no qualitative differences in our results (Supplementary Text 1).

### Stochastic evolutionary simulations

We performed Wright-Fisher simulations to describe the competition between different phenotypes in random and programmed environments. We considered discrete generations with a fixed population of bacteria. Our simulations had these steps:

1. We initialized the population in one of two ways: a. Homogeneous population, comprising a colony of a single, chosen phenotype b. Mixed population, comprising equal numbers of all possible phenotypes
2. We computed the fitness of bacteria as follows: a. In a programmed environment, we employed the sequence of stimuli *I*_1_ *→ I*_2_ *→* … *→ I*_*N*_. The fitness of each phenotype was the time-average of the fitness *ϕ* (*t*) when all the N signals were elicited once:

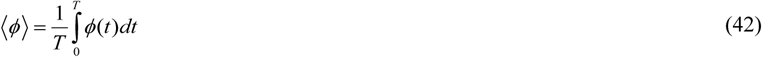

Here, *T* was chosen to be the time when the last signal faded away. b. In a random environment, the signals were elicited in a random sequence. Thus, N^N^ signal sequences were possible, allowing for the signals to repeat. The fitness of each phenotype was then the mean of its time-averaged fitness estimated separately for each of the N^N^ possible sequences:

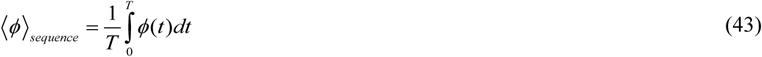

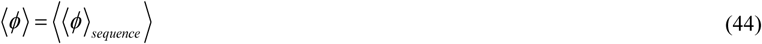
3. We next estimated ‘control’ fitness, measuring the reduction in fitness in the absence of any response, using:

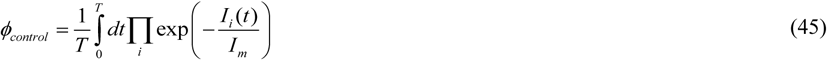

This has the same expression as *ϕ*_*i*_, but without the *f*_*b*_ term.
4. Fitness selection happens on the bacteria in a generation. For each bacterium, we examined whether the fitness *ϕ* was larger than *ϕ*_*control*_ +(1−*ϕ*_*control*_) *× r*, where *r* ∈[0,1] was a random number from a uniform distribution. The latter choice accounted for any stochastic variations in environmental factors and associated selection forces. If ⟨*ϕ*⟩ was larger, the bacterium survived. Else, it was removed.
5. From the survivors, we randomly selected some and duplicated them to replace lost bacteria and maintain the population constant.
6. We mutated the resulting bacteria. In our simulations, a mutation toggled a potential crosstalk interaction between on and off. For instance, for a bacterium with crosstalk between *HK*_*i*_ and *RR*_*j*_, mutation would turn the corresponding *k*_*f,ij, phtrf*_ and *k*_*f,ij, phtse*_ from γ×10^−3^ nM^-1^s^-1^ to 0. Every bacterium was checked for the possibility of mutation with probability μ at each of the 2^N(N-1)^ crosstalk interactions possible.
7. We repeated the above procedure from step 4.

One generation in our simulation time frame was typically *T*=N×500 s, with N signals elicited in each generation. This made sure that all the TCSs could be triggered in principle. We performed simulations over a large number of generations and over 50 realizations for each parameter setting, which ensured reliable statistics.

## Codes and data availability

The MATLAB codes used to estimate the fitness values, perform Wright-Fisher simulations, and the codon and amino acid sequence files, domain information, alignment files, and the raw data for the resulting phylogenetic trees employed for evolution analyses are available at the GitHub repository https://github.com/vembha/TCS_crosstalk_evolution.

## Acknowledgements

We thank Sandhya Visweswariah and Supreet Saini for comments and Gaurav Sankhe for inputs and discussions.

## Author Contributions

BV, AVP, NMD designed the problem. BV, AVP developed the models and codes and performed calculations. BV, AVP, DKS, NMD, analyzed the data. BV, NMD wrote the paper. BV, AVP, DKS, NMD edited the paper.

## Competing Interests

The authors declare that they do not have any competing interests.

## Notes

### Competing Interest Statement

The authors have declared no competing interest.

