## Supplementary information for "An evolutionary paradigm favoring crosstalk between bacterial two-component signaling systems"

**This PDF file includes:**

Supplementary Text 1 to 2

Supplementary Figures 1 to 7

Supplementary Tables 1 to 2

References

#### SUPPLEMENTARY TEXT 1: Alternative fitness formulation

To test the robustness of our predictions to the details of the fitness formulation, we constructed an alternative fitness function and evaluated its implications:

$$\phi_i = 1 - \frac{I_i(t)}{I_m + I_i(t)} \times \frac{K_1}{K_1 + (RR_i^*)^2} \quad \text{(S1)}$$

In the absence of the input signal $I_i$, the fitness is the maximum, i.e., one. The fitness decreases with increase in $I_i$ in a saturable manner with the half-saturation concentration $I_m$. The fitness recovers with the response, which is proportional to the fraction of free promoters $f_f$. The expression for $f_f$ derived earlier (Eq. (18) in Methods) is employed above. This alternative formulation is used to estimate the fitness of various phenotypes as in Fig. 1. The selection coefficients of all the phenotypes for N=2 in programmed and random environments, respectively, estimated for several values of $\gamma$ are in Supplementary Fig. 2a and Supplementary Fig. 2b. Phenotype 2, with one-way crosstalk mirroring the signal sequence, had the highest fitness in a programmed environment, whereas phenotype 1, without any crosstalk, had the highest fitness in a random environment, indicating the robustness of our results to the fitness formulation.

#### SUPPLEMENTARY TEXT 2: $K_A/K_S$ analysis

Previous studies have argued that strong evolutionary pressure drives the diversification of TCS genes to ensure specificity following the creation of a new TCS by gene duplication (1, 2). We hypothesized that this pressure must be substantially lower, if at all, for TCSs between which crosstalk would be favored because of adaptation to a programmed environment. To test this hypothesis, following previous studies(1), we analyzed the sequences of the TCS genes of *M. tuberculosis* and estimated the ratio of non-synonymous to synonymous mutations ($K_A/K_S$). $K_A/K_S$ analysis has been employed previously to analyze the propensity of crosstalk between TCSs (1). Here, $K_S$ is the number of synonymous mutations per synonymous site and $K_A$ is the number of non-synonymous mutations per non-synonymous site. In general, if $K_A/K_S<1$, then selection pressure is thought to conserve protein sequences, whereas if $K_A/K_S>1$, the pressure is to diversify the genes (1, 3). For crosstalk, the binding regions of the noncognate HK and RR pairs must not be significantly different from those of their corresponding cognate pairs. Evolutionary pressure must therefore act to conserve these sequences post gene duplication. In other words, $K_A/K_S$ for TCSs that crosstalk must be smaller than for specific TCSs.

We downloaded both the nucleotide and amino acid sequences of the TCSs of *M. tuberculosis* from the National Center for Biotechnology Information's (NCBI) GenBank database (4). We identified the kinase domain of each HK and the receiver domain of each RR using InterPro (5). These are the domains involved in the binding of HKs with RRs, required for phosphotransfer. We next aligned the nucleotide sequences of the domains, guided by their corresponding amino acid alignments, using the online tool Clustal Omega (6) (Supplementary Fig. 6a, Supplementary Fig. 6b) and evaluated the sequence similarity of the HK and RR domains (Supplementary Fig. 6c, Supplementary Fig. 6d).

Next, we identified TCSs for which $K_A/K_S$ could be estimated. Entities that share ancestry in gene duplication events have the possibility of crosstalk (7). Using whole protein amino acid sequences, we constructed phylogenetic trees separately for HKs (Supplementary Fig. 7a) and RRs (Supplementary Fig. 7b) following the maximum likelihood method with 500 bootstrap replicates using the MEGA (version 7) software package (8). We aligned the domain sequences of HKs and RRs separately. We considered HK-RR pairs connected within two levels of ancestry for $K_A/K_S$ estimation. We assumed that the corresponding TCSs had arisen from gene duplication events. For TCSs farther away in the trees, acquisition by horizontal gene transfer is more difficult to rule out (7). The $K_A/K_S$ ratios were estimated using the MEGA (version 7) software package (8). We calculated *p* values using a one-tailed paired *t*-test.

TCSs of *M. tuberculosis* have been classified into two major groups based on their homology (9). The first group, the NarL family, consists of Nar, Dev, and Pdta TCSs. The second group, the OmpR family, contains the rest. Our trees were consistent with this classification. (We neglected the atypical TCS Rv0600c-Rv0601c-TcrA, which does not undergo autophosphorylation (10).) From the trees, we identified HKs and RRs connected within two levels of ancestry. This yielded the following related HKs (Supplementary Fig. 7a): 1) NarS and DevS/DosT (green node); 2) MprB, PrrB, and SenX3 (blue node); and 3) PhoR, TrcS, and TcrY (yellow node). Similarly, the following related RRs emerged (Supplementary Fig. 7b): 1) PdtaR, NarL, and DevR (green node); 2) MprA, PrrA, and KdpE (blue node); and 3) PhoP, TrcR, and TcrX (yellow node). Note that the HKs and RRs in nodes of the same color belong largely to the same TCSs. For instance, the yellow node has the PhoR-PhoP, TrcS-TrcR, and TcrY-TcrX TCSs in both the trees. This gave us further confidence in their possible connectedness through gene duplication. The kinase domain has not been annotated for the NarS gene(5). We thus could not analyze the green node. We analyzed sequence data from the other two nodes.

We recall from the *in vitro* data (Main text, Fig. 4a) that in the blue nodes, PhoR-PhoP and TcrY-TcrX crosstalk, whereas, in the yellow nodes, MprB-MprA and PrrB-PrrA crosstalk. The other pairs do not. We estimated the $K_A/K_S$ ratios for all the HK pairs and the RR pairs in the yellow and blue nodes (Supplementary Fig. 7c, Supplementary Table 2). The ratios were 0.59±0.18 for the HKs and 0.57±0.09 for the RRs, indicating no significant overall difference in evolution between the HKs and RRs. Because the $K_A/K_S$ ratios can vary between the nodes, to assess the differences between the $K_A/K_S$ ratios of TCSs engaged in crosstalk and not, we performed the one-tailed paired *t*-test (paired by their nodes in the phylogenetic tree, since the genes are assumed to have been duplicated) for the HKs and RRs separately. Interestingly, we found that the ratios for the RRs engaged in crosstalk (PrrA-MprA and TcrX-PhoP) were significantly lower than for those that did not ($p$=0.035). The difference was not significant for the HKs ($p$=0.433). The implication is that the binding domains in the RRs that crosstalk were under much less evolutionary pressure to diversify than those that were specific. The HKs were not under similar pressure. Note that crosstalk is possible if either the HKs or the RRs have their binding domains conserved. It is only if both diversify that specificity results. For instance, if $HK_1$ is similar to $HK_2$ but $RR_1$ is removed from $RR_2$, $HK_1$ will still be able to crosstalk to $RR_2$, because $HK_2$ will continue to exhibit its cognate interaction with $RR_2$. $HK_1$ and $HK_2$ must also become dissimilar for the two TCSs to become insulated. We may infer thus that TCSs that crosstalk may be under less evolutionary pressure to diversify post gene duplication than TCSs that are specific.

### SUPPLEMENTARY FIGURES

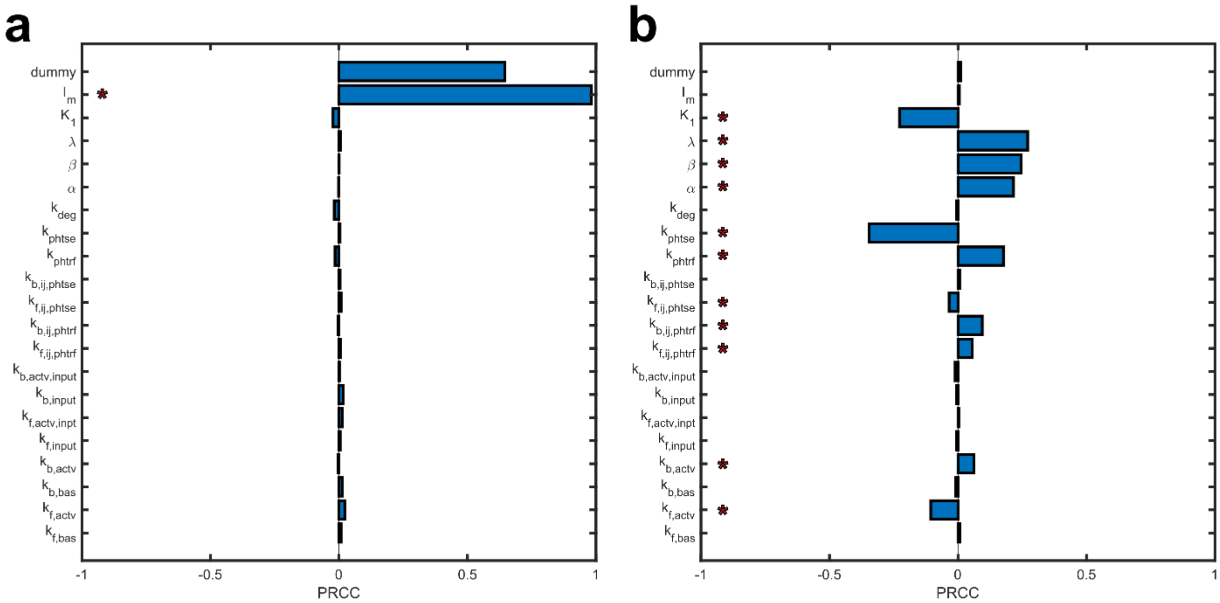


**Supplementary Fig. 1 Sensitivity analyses of the model.** (**a**) Partial rank correlation coefficients (PRCCs) (11) indicating the sensitivity of the fitness of a single TCS to model parameters. (**b**) Sensitivity of the selection coefficient of phenotype 2 (with crosstalk between $HK_1$-$RR_2$) for a bacterium with N=2 TCSs in a programmed environment. The red asterisks indicate $p<0.01$. The fitness was thus significantly sensitive to the signal strength $I_m$. The selection coefficient was sensitive to phosphotransfer ($k_{phtrf}$) and phosphatase ($k_{phtse}$) rates, as well as the parameters that affect events post $RR^*$- DNA binding $(\alpha, \beta, \lambda)$. This sensitivity influences our predictions quantitatively and not qualitatively, leaving our inferences robust.

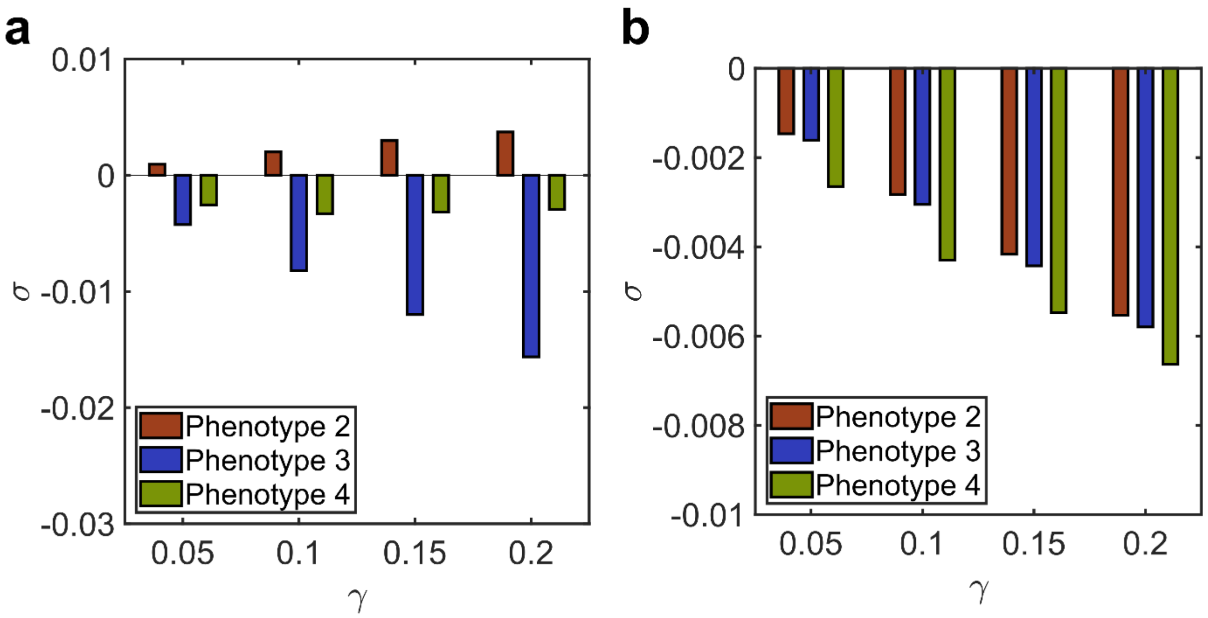


**Supplementary Fig. 2 Selection coefficient with alternative fitness formulation for N=2.** (**a**) Selection coefficient, σ, as a function of crosstalk strength, γ, when signal 2 follows signal 1. (**b**) σ as a function of γ when signals 1 and 2 follow no order. Fitness is calculated using Eq. (S1). The other details are identical to those in Fig. 1. In agreement with Fig. 1, phenotype 2, with crosstalk between $HK_1$ and $RR_2$, has the highest fitness in programmed environment while phenotype 1, with no crosstalk, has the highest fitness in a random environment.

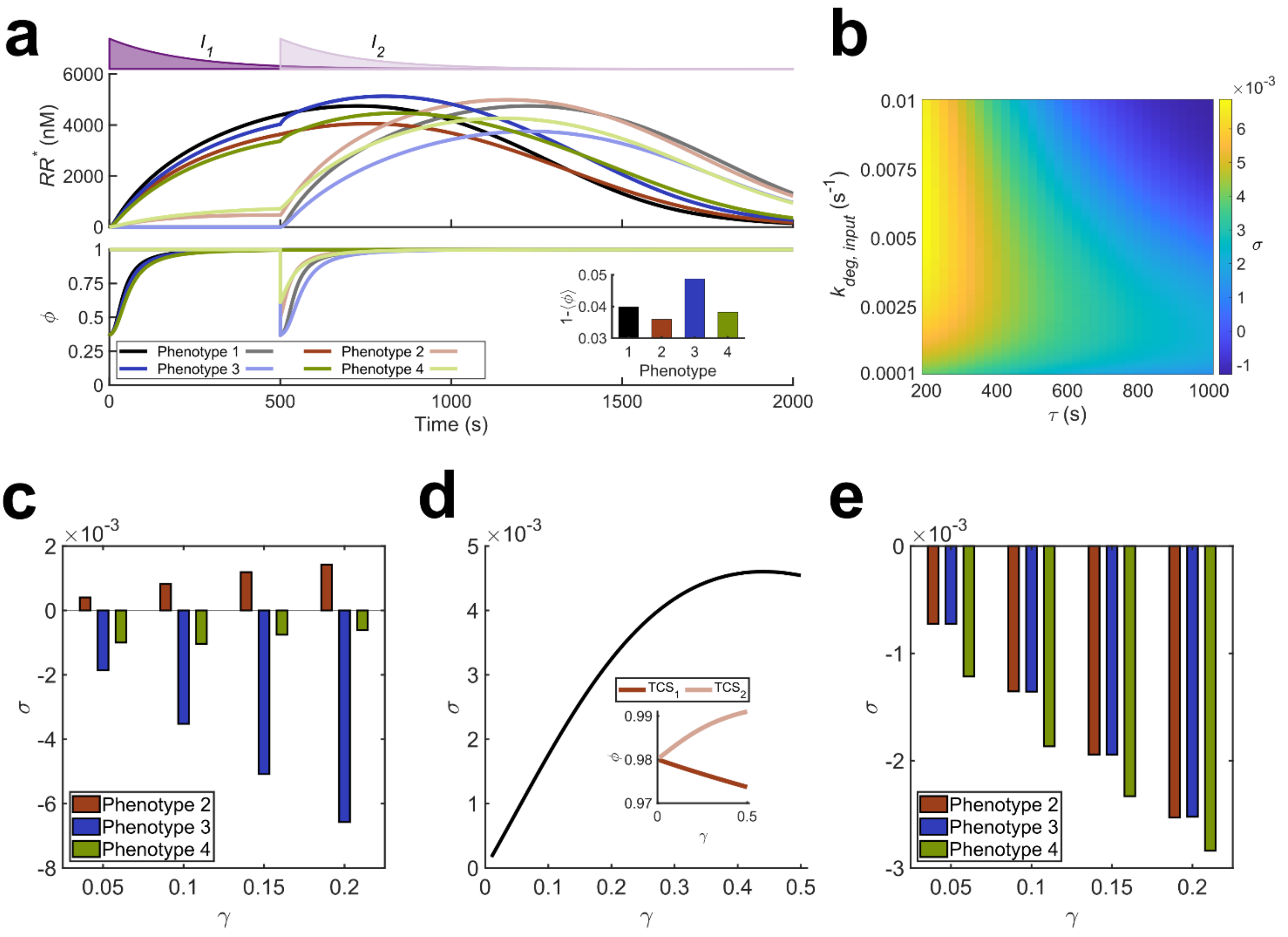


**Supplementary Fig. 3 Behavior with exponentially decaying input signal.** (**a**) Input-output behavior. The magenta filled curves on the top depict the strength of the input signals over time, with the darker curve representing $I_1$ and the lighter curve $I_2$. The signals decay exponentially with rate constant $k_{deg,input}$=4.605×10$^{-3}$ s$^{-1}$, starting from the peak of 10$^4$ nM. The signals are separated by τ=500 s. The top panel shows the concentrations of activated RRs and the bottom panel shows the associated fitness of the responses. The phenotypes are color coded and dark and light curves represent TCS1 and TCS2, respectively. Crosstalk strength is γ = 0.26. The inset shows the time-averaged fitness of the different phenotypes (description of phenotypes can be found in *Main text,* Fig. 1*A*). (**b**) Selection coefficient (σ) for phenotype 2 with varying τ and $k_{deg,input}$. σ is defined as the fitness advantage a phenotype has over the phenotype without crosstalk. (**c**) Selection coefficients in programmed environment. σ as a function of γ when signal 2 follows signal 1. (**d**) Optimal crosstalk strength. Dependence of σ on γ for phenotype 2 shows the trade-off between increasing fitness of TCS2 due to priming and decreasing fitness of TCS1 due to leakage (inset) resulting in maximum overall fitness at intermediate γ. (**e**) Selection coefficients in random environment. σ as a function of γ when signals 1 and 2 follow no order. Fitness is calculated as the mean all possible signal sequences.

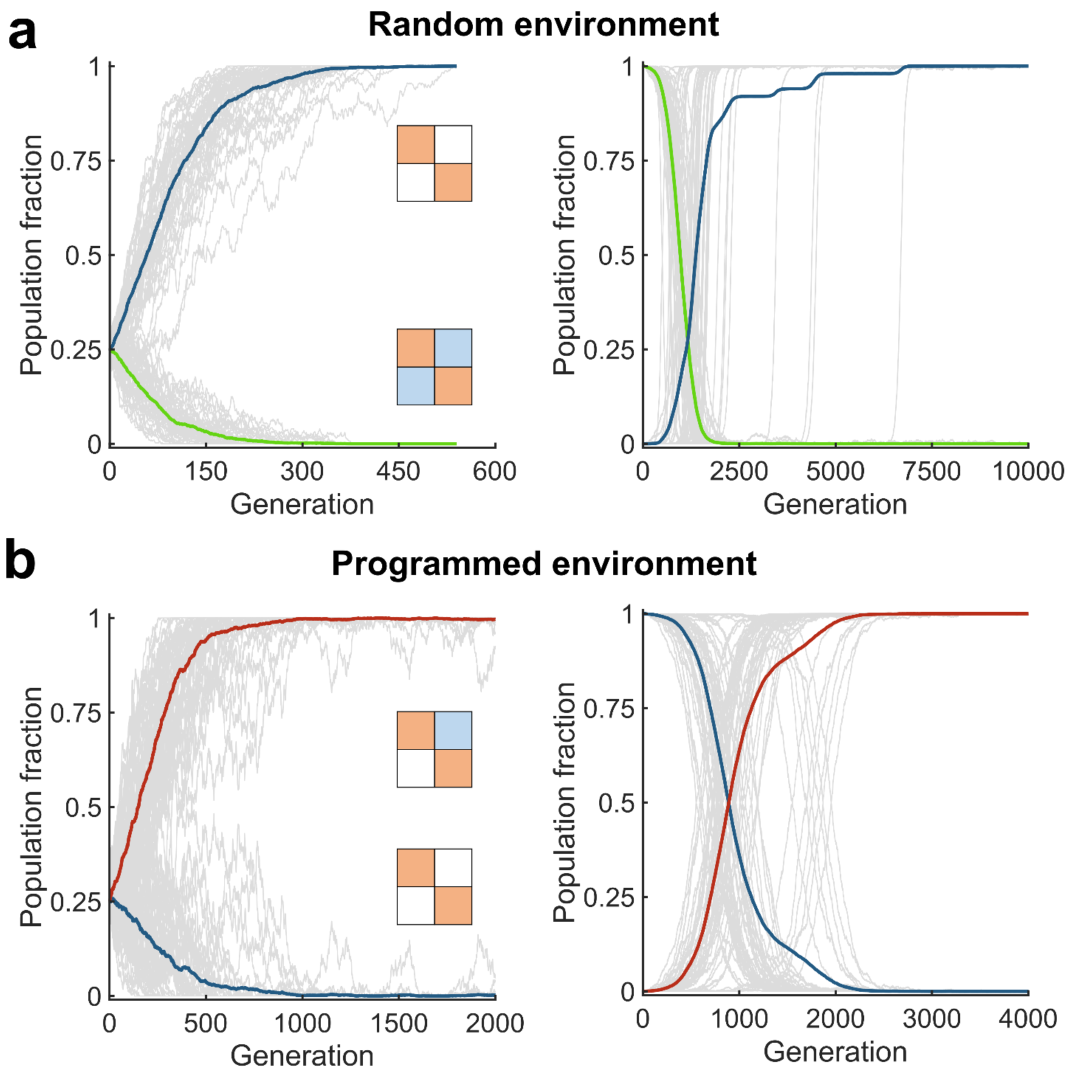


**Supplementary Fig. 4 Evolution of bacteria with N=2 TCSs.** (**a**) Evolution in a random environment. The phenotype without any crosstalk (blue) gets fixed whether the initial population is homogeneous (left) or mixed (right). The phenotype with all crosstalk interactions is also shown for comparison (green). The gray lines are trajectories of the two phenotypes in each of 50 realizations. The thick lines are means. Trajectories of all other phenotypes are not shown. (**b**) Evolution in a programmed environment. Phenotype with one-way crosstalk mirroring the signal sequence (red) dominates the population whether the initial population is homogeneous (left) or mixed (right). The crosstalk strength was set to $\gamma = 0.26$ throughout.

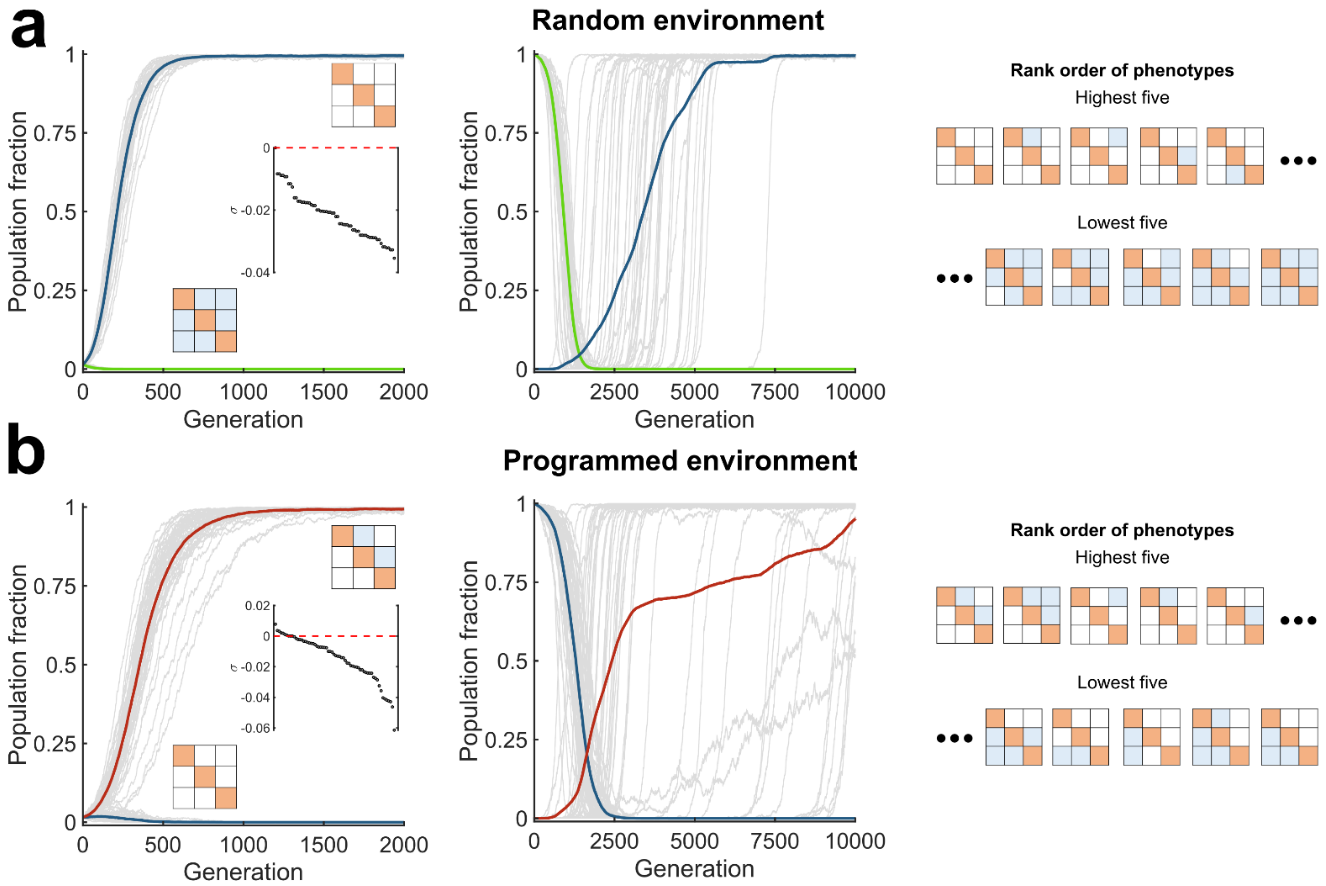


**Supplementary Fig. 5 Evolution of bacteria with N=3 TCSs.** (**a**) Evolution in a random environment. The phenotype without any crosstalk (blue) gets fixed whether the initial population is homogeneous (left) or mixed (middle). The phenotype with all crosstalk interactions is also shown for comparison (green). The gray lines are trajectories of the two phenotypes in each of 50 realizations. The thick lines are means. Trajectories of all other phenotypes are not shown. The inset (left) is the rank-ordered selection coefficient for all the phenotypes. The interaction matrices of the five most and least fit phenotypes are shown (right). (**b**) Evolution in a programmed environment. Phenotype with one-way crosstalk mirroring the signal sequence (red) dominates the population, whether the initial population is homogeneous (left) or mixed (middle). The crosstalk strength was set to γ = 0.26 throughout. The inset (left) is the rank-ordered selection coefficient for all the phenotypes. The interaction matrices of the five most and least fit phenotypes are shown (right).

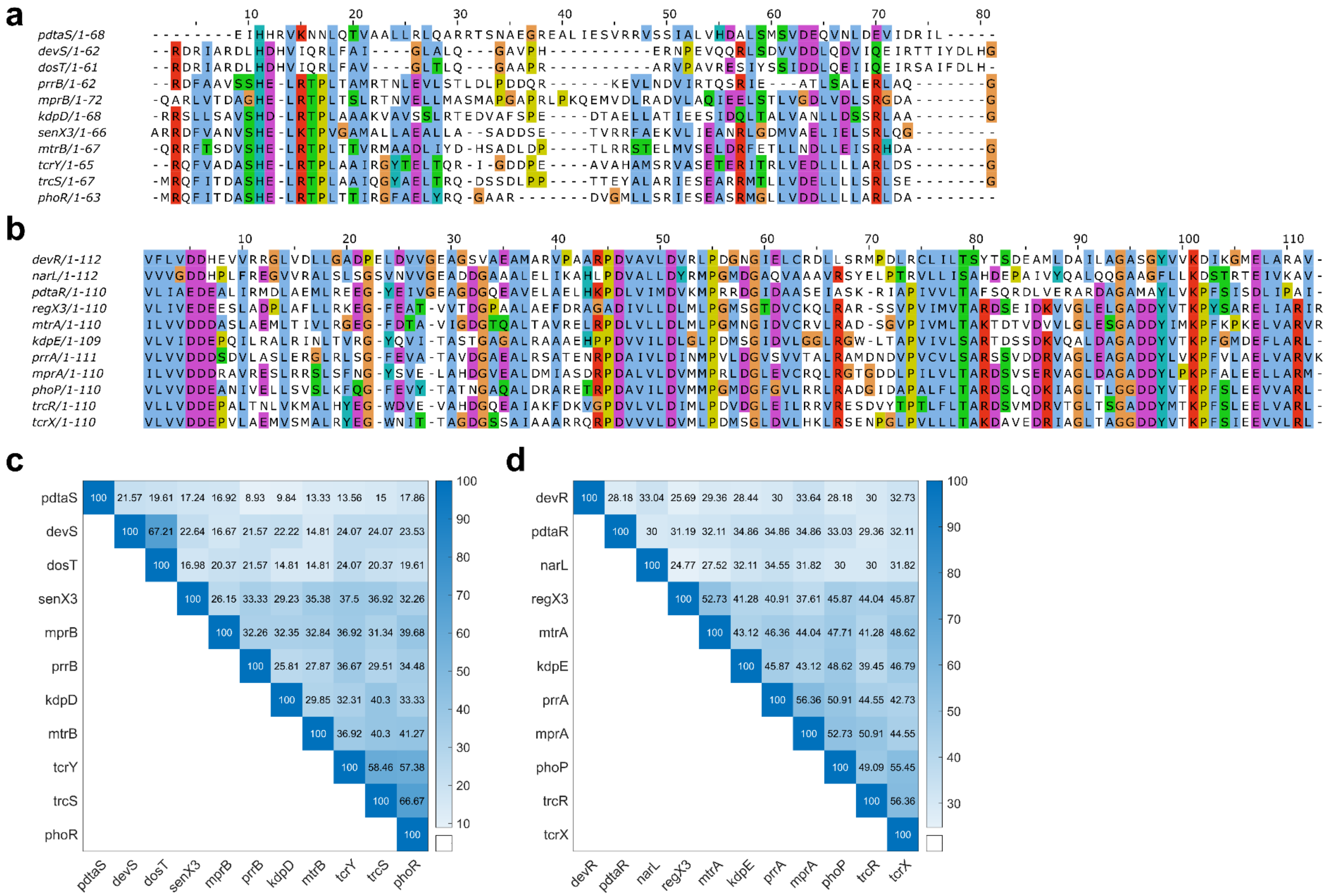


**Supplementary Fig. 6 Genetic data of domains.** Amino acid sequence alignment of (**a**) HKs and (**b**) RRs of *M. tuberculosis* used in our $K_A/K_S$ analyses. The corresponding percent similarity of these domains obtained from Clustal Omega (10) for (**c**) HKs and (**d**) RRs respectively.

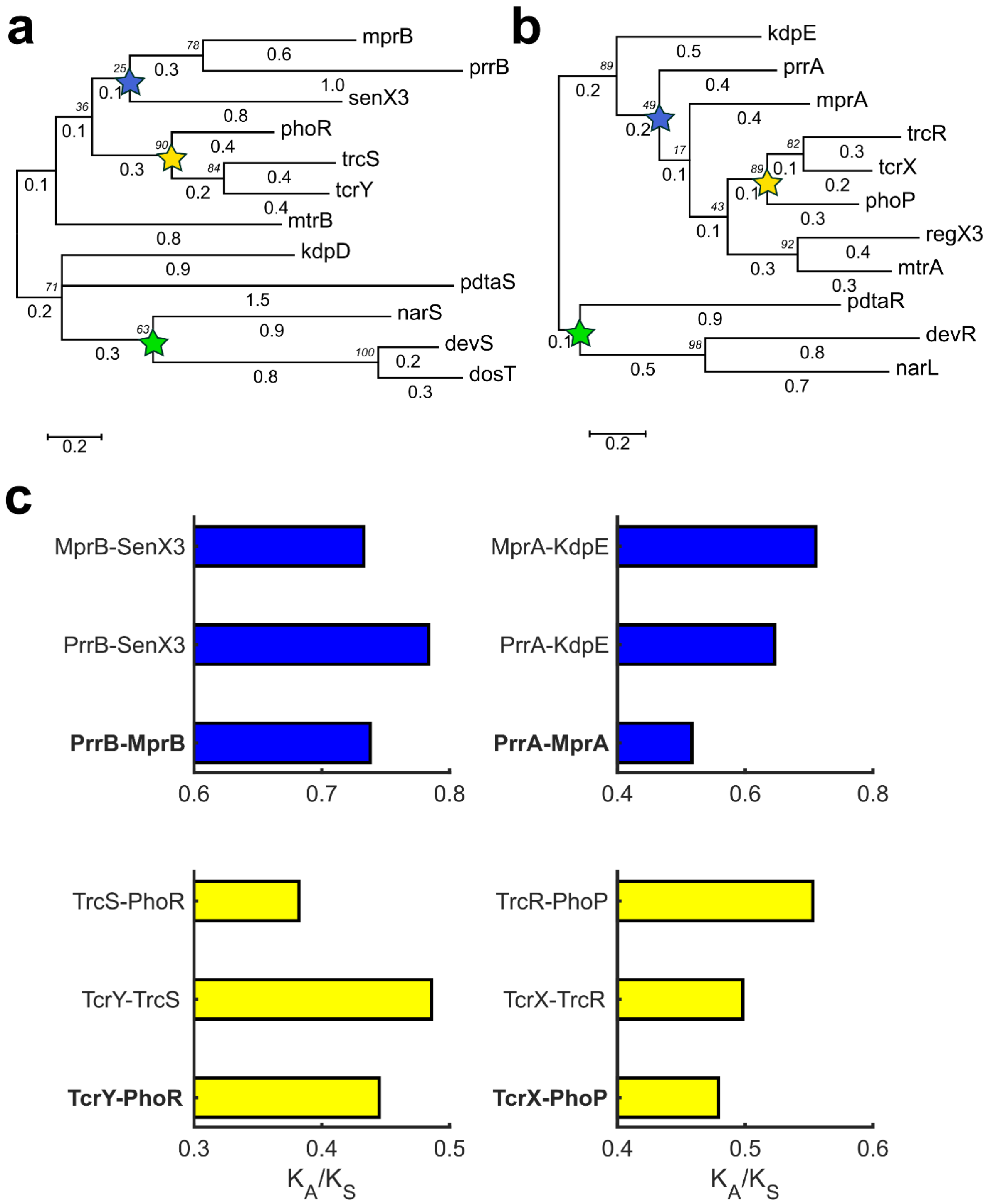


**Supplementary Fig. 7 Diversifying pressure on TCSs of *M. tuberculosis*.** Phylogenetic tree from (**a**) HK and (**b**) RR sequences. Branch lengths are presented below the branches, while the numbers in italics are the bootstrap statistics for each node. (**c**) $K_A/K_S$ ratios estimated for the binding domains of the TCS proteins in the yellow and blue nodes in (A) and (B) and are shown for HKs (left panel) and RRs (right panel) respectively; data is presented in Supplementary Table 2. Combinations marked in bold crosstalk *in vitro* (Main text, Fig. 4a).

### SUPPLEMENTARY TABLES

**Supplementary Table 1 Model parameters and initial conditions.**

| Symbol | Description | Value | Source |
|---|---|---|---|
| $P_T$ | Total promoter concentration | 100 nM | (1) |
| $k_{f,bas}$ | Basal autophosphorylation rate | $10^{-10}$ $s^{-1}$ | (1) |
| $k_{f,actv}$ | $I_iHK_i$ autophosphorylation rate | 0.1 $s^{-1}$ | (1) |
| $k_{b,bas}$ | Basal dephosphorylation rate | 0.1 $s^{-1}$ | (1) |
| $k_{b,actv}$ | $I_iHK_i^*$ dephosphorylation rate | 0.1 $s^{-1}$ | (1) |
| $k_{f,input}$ | $I_i - HK_i$ binding rate | 100 $nM^{-1}s^{-1}$ | (1) |
| $k_{f,actv,input}$ | $I_i - HK_i^*$ binding rate | 100 $nM^{-1}s^{-1}$ | (1) |
| $k_{b,input}$ | $I_iHK_i$ unbinding rate | 1000 $s^{-1}$ | (1) |
| $k_{b,actv,input}$ | $I_iHK_i^*$ unbinding rate | $10^{-6}$ $s^{-1}$ | (1) |
| $k_{f,ij,phtrf}$ | $HK_i^* - RR_j$ binding rate | $\gamma \times 10^{-3}$ $nM^{-1}s^{-1}$ | (1) |
| $k_{b,ij,phtrf}$ | $HK_i^* - RR_j$ unbinding rate, $\forall i, j$ | 1 $s^{-1}$ | (1) |
| $k_{f,ij,phtse}$ | $HK_i - RR_j^*$ binding rate | $\gamma \times 10^{-3}$ $nM^{-1}s^{-1}$ | (1) |
| $k_{b,ij,phtse}$ | $HK_i - RR_j^*$ unbinding rate, $\forall i, j$ | 1 $s^{-1}$ | (1) |
| $k_{phtrf}$ | Phosphotransfer rate | 0.055 $s^{-1}$ | - |
| $k_{phtse}$ | Cognate phosphatase activity rate | 0.01 $s^{-1}$ | - |
| | Non-cognate phosphatase activity rate | $1.67 \times 10^{-3}$ $s^{-1}$ | |
| $k_{deg}$ | Protein degradation rate | $6 \times 10^{-5}$ $s^{-1}$ | (12) |

| Symbol | Description | Value | Source |
|---|---|---|---|
| $\alpha$ | Ratio of basal and activated transcription rates | 10 | (12) |
| $\beta$ | $HK-RR$ synthesis rate | $6\times10^{-3}$ nM $s^{-1}$ | (12) |
| $\lambda$ | Ratio of $HK$ and $RR$ concentrations | 0.1 | (12) |
| $K_1$ | Dissociation constant for $(RR_j^*)^2 P_j$ | $5\times10^{5}$ $nM^2$ | (12) |
| $\mu$ | Mutation rate | $10^{-5}$ per crosstalk interaction, per generation | Assumed |
| $k_{deg,input}$ | Input degradation rate | 0 (for step input)<br>$4.605\times10^{-3}$ $s^{-1}$ (for exponentially decaying input) | Assumed |
| $\gamma$ | Fold change in $k_{f,ij,phtrf}$ and $k_{f,ij,phtse}$ | 1 if $i=j$, else varied | - |
| $HK(0)$ | Initial concentration of HKs | 100 nM | (1) |
| $RR(0)$ | Initial concentration of RRs | 1000 nM | (1) |
| $I_m$ | Peak stimulus strength | $10^4$ nM | (1) |

**Supplementary Table 2 $K_A/K_S$ ratios.** The $K_A/K_S$ ratios for the HKs and RRs identified in the colored nodes of the phylogenetic trees in the Supplementary Fig. 7. The entities highlighted in the respective color are the ones predicted to crosstalk (Main text, Fig. 4a).

| **Node** | ***$HK_1$*** | ***$HK_2$*** | **$K_A/K_S$** | ***$RR_1$*** | ***$RR_2$*** | **$K_A/K_S$** |
|---|---|---|---|---|---|---|
| **Blue** | PrrB | MprB | 0.7379 | PrrA | MprA | 0.5165 |
| | PrrB | SenX3 | 0.7836 | PrrA | KdpE | 0.6461 |
| | MprB | SenX3 | 0.7328 | MprA | KdpE | 0.7104 |
| **Yellow** | TcrY | PhoR | 0.4450 | TcrX | PhoP | 0.4791 |
| | TcrY | TrcS | 0.4858 | TcrX | TrcR | 0.4981 |
| | TrcS | PhoR | 0.3821 | TrcR | PhoP | 0.5529 |
